# The KAT2 Acetyltransferase PCAF-1 Couples Stress-Responsive Transcription to Proteostasis and Longevity

**DOI:** 10.64898/2026.11.10.765553

**Authors:** Avinash Sharma, Khairun Nisaa, Lina Blanco Rodríguez, Jeremy Tachiki, Erica Dinneen, Mai Huong To, Carlos G. Silva-García

## Abstract

Aging erodes the capacity of cells to sense stress and maintain a functional proteome. Lysine acetyltransferases can link stress signals to gene expression, but whether boosting the activity of a single acetyltransferase can enhance stress responses to protect the aging proteome and extend healthspan remains unknown. Here we show that increasing the dosage of PCAF-1, the sole KAT2 family acetyltransferase in *Caenorhabditis elegans*, is sufficient to extend lifespan, preserve healthspan, and reprogram stress-responsive transcription. This program is conserved; mammalian orthologs of PCAF-1-induced genes are enriched among transcripts that decline upon KAT2A and KAT2B depletion. PCAF-1 strengthens inducible stress responses, attenuates the age-dependent decline in proteasomal degradation, and limits protein aggregation in *C. elegans* models of human neurodegenerative diseases, while preserving motility and survival. Mechanistically, PCAF-1 drives DAF-16/FOXO activation, and DAF-16 is required for both PCAF-1-dependent longevity and for protection against polyglutamine aggregation. These findings reveal a previously unrecognized mechanism by which KAT2 acetyltransferases promote longevity, positioning KAT2s as potential tunable regulators of proteostasis and healthspan during aging.

## INTRODUCTION

Aging is accompanied by the accumulation of molecular damage and a progressive decline in the capacity to sense and respond to it^1^. As this adaptive capacity weakens, the systems that maintain protein quality and mount protective stress responses become less effective. Because many of these responses depend on changes in gene expression and transcriptional regulation, chromatin-modifying enzymes are well positioned to influence how organisms preserve function during aging^2^. Lysine acetyltransferases (KATs), many of which were first characterized as histone acetyltransferases^3^, are particularly relevant in this context because they can act on both chromatin and non-histone substrates^4^, including transcription factors^5^, allowing them to connect physiological signals with stress-responsive transcription and cellular maintenance.

In *C. elegans*, the MYST-family acetyltransferase complex MYS-1–TRR-1 promotes stress resistance and longevity^6^, and the CBP/p300 ortholog CBP-1, which belongs to the KAT3 family of lysine acetyltransferases, is required for proteostasis and lifespan extension during mitochondrial stress^7^. However, acetyltransferases can also counteract protective programs. CBP-1 restrains the heat shock response in *C. elegans*^8^, whereas the KAT2-family member GCN5 acetylates and inhibits TFEB, thereby suppressing autophagy and lysosome biogenesis in mammalian cells and *Drosophila* ^9^. Furthermore, partial reduction of Gcn5 in yeast extends lifespan^10^. Thus, the effects on proteostasis and aging depend on the enzyme and context and are largely defined by loss-of-function studies under specific stress conditions. Whether KAT2-family enzymes promote or oppose stress resilience during aging, and whether increasing their activity is sufficient to preserve organismal function and longevity, remains unknown.

PCAF-1 is the sole *C. elegans* member of the conserved KAT2 family of lysine acetyltransferases, represented in mammals by KAT2A/GCN5 and KAT2B/PCAF. KAT2-family proteins are evolutionarily conserved transcriptional regulators that function within multiprotein coactivator complexes, including SAGA (Spt–Ada–Gcn5 acetyltransferase) and ATAC (Ada-two-A-containing), which provide distinct molecular contexts for their acetyltransferase activity^11^. As the sole KAT2-family member in *C. elegans*, PCAF-1 offers an opportunity to investigate the physiological consequences of KAT2 activity without the paralog redundancy found in mammals.

Beyond transcriptional activation, KAT2-family acetyltransferases act as metabolic sensors that couple nutrient and stress signals to gene expression. In yeast, the AMPK orthologue Snf1 promotes recruitment of Gcn5 to a glucose-responsive promoter, along with increased histone H3K9 and H3K14 acetylation and chromatin remodeling^12^. In human cells, AMPK activation also stimulates KAT2B-dependent acetylation^13^. In mammals, KAT2B is required for pancreatic beta-cell adaptation to metabolic challenge, where it sustains expression of endoplasmic-reticulum unfolded protein response genes^14^. Together, these findings place KAT2 acetyltransferases at the interface of nutrient signaling and stress adaptation in evolutionarily distant organisms. We recently identified PCAF-1 as a pro-longevity factor: it is required for the extended lifespan of mutants lacking components of COMPASS (complex of proteins associated with Set1), and increasing its dosage alone is sufficient to extend wild-type lifespan^15^. Yet how PCAF-1 preserves cellular function during aging, and which factors mediate this effect, remained undefined.

Here, we show that increased PCAF-1/KAT2 expression reprograms stress-responsive transcription together with programs governing metabolic regulation and protein quality control, with these effects conserved in mammalian cells. Elevated PCAF-1 enhances UPR^ER^ inducibility, limits the age-associated accumulation of a proteasome substrate, and protects against polyglutamine and α-synuclein proteotoxicity. PCAF-1 further promotes DAF-16/FOXO nuclear localization in a manner sensitive to acetyltransferase inhibition, and DAF-16 is required for both PCAF-1-mediated longevity and protection against toxic aggregates. Finally, HLH-30/TFEB and AAK-2/AMPK are additionally required for lifespan extension, and AMPK acts downstream of PCAF-1, potentially through the creatine-kinase-like effector ARGK-1. Together, these findings identify PCAF-1 as a potential pro-longevity acetyltransferase that couples stress-responsive transcription to proteostasis, promoting resilience and longevity during aging through a conserved AMPK, DAF-16/FOXO, and HLH-30/TFEB dependent program.

## RESULTS

### PCAF-1 promotes stress-adaptive transcription and preserves organismal health during aging

Previously, we showed that increased expression of the KAT2 lysine acetyltransferase PCAF-1 drastically extends lifespan in *C. elegans*^15^. However, the molecular mechanisms by which PCAF-1 extends lifespan remain unknown. To investigate this, we performed RNA sequencing (RNA-seq) of wild-type (N2) animals and a transgenic line constitutively expressing *pcaf-1* (*pcaf-1 Tg*). Principal component analysis shows a clear separation between wild-type and *pcaf-1 Tg* samples **(Fig. 1a)**. We identified 3,068 differentially expressed genes, including 2,334 upregulated and 734 downregulated genes in *pcaf-1 Tg* animals **(Fig. 1b)**. Complete gene-level differential-expression results are provided in **Supplementary Data 1**. KEGG overrepresentation analysis of these genes shows that genes induced in *pcaf-1 Tg* animals are enriched for pathways involved in the integrated stress response, xenobiotic and glutathione metabolism, amino-acid and fatty-acid metabolism, and longevity regulation **(Fig. 1c)**. In contrast, genes repressed in *pcaf-1 Tg* animals are enriched for ribosome biogenesis, RNA polymerase, DNA replication and RNA degradation, as well as ubiquitin-mediated proteolysis and proteasome-associated pathways **(Fig. 1d)**. Several of these features, particularly stress-response gene expression and reduced representation of ribosome-biogenesis machinery, have been associated with longevity in *C. elegans*^16–18^.

**Fig. 1.**
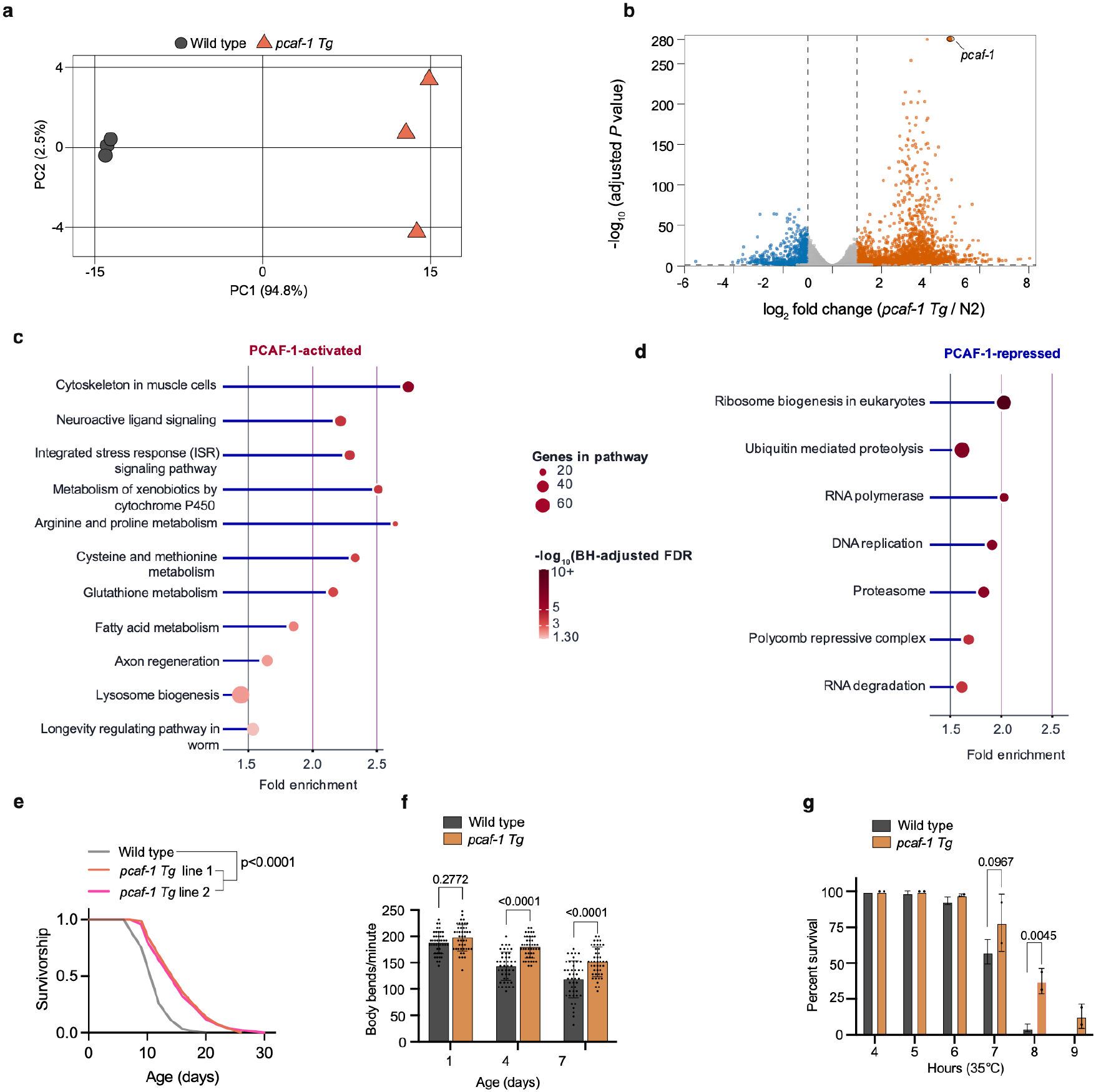
PCAF-1 promotes stress-adaptive transcription and preserves organismal function during aging. **a,** Principal-component analysis (PCA) of RNA-seq data from wild-type and *pcaf-1* transgenic (*pcaf-1 Tg*) day-1 adult animals. Three independent biological replicates per genotype. PC1 and PC2 account for 94.8% and 2.5% of the total variance, respectively. **b,** Volcano plot showing differential gene expression in *pcaf-1 Tg* animals relative to wild-type. A total of 3,068 differentially expressed genes were identified, comprising 2,334 upregulated and 734 downregulated genes (adjusted *P* < 0.05 and |log₂ fold change| ≥ 1). The x axis represents log₂ fold change, and the y axis represents −log₁₀(adjusted *P* value). **c,d,** KEGG pathway over-representation analysis of genes upregulated (**c**) and downregulated (**d**) in *pcaf-1 Tg* animals. To capture pathways containing genes with smaller but significant expression changes, gene sets were defined using adjusted *P* < 0.05 and |log₂ fold change| ≥ log₂ (1.2). Enrichment was assessed using a one-sided hypergeometric test with Benjamini–Hochberg correction for multiple testing. Selected significantly enriched pathways (FDR < 0.05) are shown. Dot size represents the number of genes associated with each pathway, dot color represents −log₁₀-adjusted FDR, and the x-axis indicates fold enrichment. **e,** Survival curves of wild-type animals and two independently generated *pcaf-1 Tg* lines (lines 1 and 2). Both transgenic lines exhibited significantly extended lifespan compared with N2 animals (log-rank (Mantel–Cox) test, *P* < 0.0001 for each comparison). **f,** Locomotor activity of wild-type and *pcaf-1 Tg* (line 1) animals on days 1, 4 and 7 of adulthood, measured as body bends per minute. Approximately 14–20 animals per genotype and age were examined in each of three independent biological experiments. Bars represent mean ± s.d., with individual animals shown as dots. Statistical significance was assessed using ordinary two-way ANOVA followed by Bonferroni’s multiple-comparisons test. **g,** Survival of day-1 adult wild-type and *pcaf-1 Tg* (line 1) animals exposed to acute heat stress at 35 °C. Percent survival was determined at the indicated time points over two independent biological experiments, with approximately 60 animals per genotype per experiment. Bars represent mean ± s.d. Statistical significance was assessed using two-way ANOVA followed by Bonferroni’s multiple-comparisons test. *pcaf-1 Tg* animals showed significantly greater survival at 8 h (adjusted *P* = 0.0045).

Molecular function analysis reveals enrichment of protein kinase activity, transporter activity and oxidoreductase activity among induced genes, whereas chromatin binding, histone-modifying activity and translation factor activity are enriched among repressed genes **(Extended Data Fig. 1b)**. Because several major longevity pathways in *C. elegans* are regulated through phosphorylation-dependent signaling^19^, we specifically examined whether kinase-associated genes are altered by PCAF-1 overexpression. Among genes annotated with protein kinase activity (GO:0004672), 127 are significantly induced by at least twofold, whereas 13 are significantly repressed to the same extent (**Extended Data Fig. 2**). The induced kinase-associated genes increase consistently across the three independent *pcaf-1 Tg* RNA-seq replicates, indicating that this functional class constitutes a substantial component of the PCAF-1-dependent transcriptional response. Consistent with a potential connection to longevity signaling, the RNA-seq dataset also shows significant induction of *unc-43* and *pkc-2*, two kinases previously linked to DAF-16-dependent lifespan regulation^20,21^. Overall, our data suggest that constitutive expression of PCAF-1 induces a transcriptional drift that particularly involves pathways associated with stress response and longevity.

We next examined whether increased PCAF-1 dosage preserved organismal function during aging. We first confirmed that two independently generated *pcaf-1 Tg* lines carrying the same transgene exhibit extended lifespan compared with wild-type animals **(Fig. 1e)**. To assess age-dependent functional decline, we measured locomotor activity during adulthood. Body-bend rates are comparable between wild-type and *pcaf-1 Tg* animals on day 1 but are significantly higher in transgenic animals on days 4 and 7 **(Fig. 1f)**. We also examined resistance to acute heat stress in day-1 adults. Following exposure to 35 °C, *pcaf-1 Tg* animals show greater survival than wild-type animals, with a significant difference at 8 h **(Fig. 1g)**. Taken together, our findings show that elevated PCAF-1 reprograms gene expression toward stress resistance, metabolic adaptation, and longevity, extending lifespan, preserving motility and strengthening resistance to acute heat stress.

### PCAF-1 activates a transcriptional program conserved across mammalian KAT2 models

Having identified transcriptional and physiological changes associated with increased PCAF-1 dosage, we next asked whether the PCAF-1-dependent transcriptional response is evolutionarily conserved. We first examined the conservation of PCAF-1 by comparing amino acid sequences. Phylogenetic analysis identifies PCAF-1 as the single *C. elegans* representative of the conserved KAT2 acetyltransferase family, which includes vertebrate paralogues KAT2A and KAT2B, also known as GCN5 and PCAF, respectively **(Fig. 2a and Extended Data Fig. 3)**. Comparison of protein domain architecture reveals conservation of the PCAF homology region, acetyltransferase domain and C-terminal bromodomain across metazoan KAT2 proteins **(Fig. 2b)**. At 767 amino acids, PCAF-1 is comparable in length to its approximately 800-amino-acid mammalian counterparts but substantially longer than the 439-amino-acid yeast Gcn5 protein. Multiple-sequence alignment further shows substantial amino-acid conservation within the catalytic acetyltransferase region, with 60.3% and 63.1% sequence identity between *C. elegans* PCAF-1 and human KAT2A and KAT2B, respectively **(Extended Data Fig. 4)**, supporting the evolutionary relationship between worm PCAF-1 and mammalian KAT2 proteins.

**Fig. 2.**
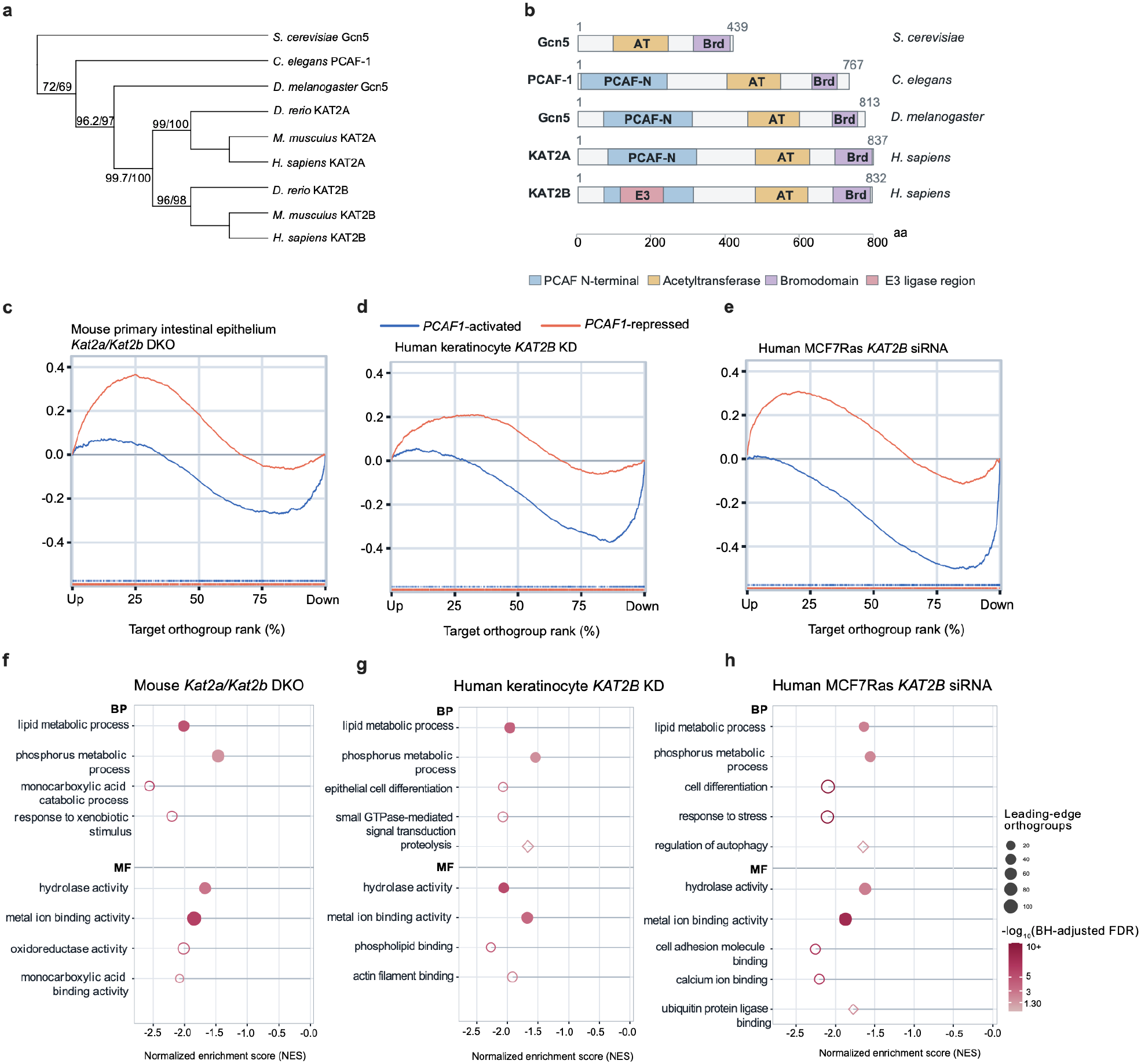
PCAF-1 regulates a transcriptional program conserved across mammalian KAT2 models. **a,** Phylogenetic tree showing nine representative GCN5/PCAF/KAT2-family acetyltransferases, derived from a maximum-likelihood analysis of 22 full-length protein sequences. Node labels indicate SH-aLRT/ultrafast-bootstrap support percentages based on 1,000 replicates each. Branch lengths are not proportional to evolutionary distance. **b,** Comparison of protein domain architecture among *C. elegans* PCAF-1, human KAT2A and KAT2B, *Drosophila melanogaster* Gcn5 and *Saccharomyces cerevisiae* Gcn5. Domains shown include the PCAF N-terminal region, acetyltransferase domain (AT) and bromodomain (Brd); an E3 ligase region is also shown for human KAT2B. Protein lengths are indicated in amino acids (aa). **c–e,** Gene set enrichment analysis (GSEA) of PCAF-1-activated and PCAF-1-repressed transcriptional programs across mammalian orthogroups ranked by their response to KAT2 depletion. Analyses include mouse primary intestinal epithelial cells lacking *Kat2a/Kat2b* (**c**; GEO: GSE235789), human keratinocytes following *KAT2B* knockdown (**d**; GEO: GSE215244) and human MCF7Ras cells treated with *KAT2B* siRNA (**e**; GEO: GSE195761). Worm gene sets were defined using adjusted *P* < 0.05 and |log₂ fold change| ≥ 0.263 and mapped to mammalian orthogroups. The x-axis represents orthogroup rank, ordered from increased to decreased expression following KAT2 loss, and the y-axis shows the running enrichment score. Tick marks indicate program-member positions. Enrichment was assessed using 100,000 permutations, with Benjamini–Hochberg correction across the six program-by-dataset comparisons (FDR < 0.05). **f–h,** Rank-based GSEA of Gene Ontology (GO) biological process (BP) and molecular function (MF) categories within the PCAF-1-activated program in mouse *Kat2a/Kat2b* double-knockout (**f**), human keratinocyte *KAT2B* knockdown (**g**) and human MCF7Ras *KAT2B* siRNA (**h**) datasets. GO-defined subsets of PCAF-1-activated orthogroups were tested against the complete ranked mammalian orthogroup datasets. Selected significantly enriched categories with negative normalized enrichment scores (NES) and Benjamini–Hochberg-adjusted FDR < 0.05 are shown. Multiple-testing correction was performed separately for each dataset and GO ontology. The x-axis represents NES, symbol size indicates the number of leading-edge orthogroups, and color represents −log₁₀-adjusted FDR, visually capped at 10. Filled circles indicate categories significantly enriched across all three mammalian datasets, open circles indicate dataset-specific categories, and open diamonds identify selected context-specific categories discussed in the results.

To determine whether this conservation extends to transcriptional regulation, we mapped PCAF-1-responsive genes to mammalian orthogroups and compared their expression patterns with published transcriptomic datasets from mouse intestinal epithelial cells lacking both *Kat2a* and *Kat2b*, human keratinocytes depleted of *KAT2B*, and human MCF7Ras cells treated with *KAT2B* siRNA^22–24^. Gene set enrichment analysis reveals that orthogroups induced in *pcaf-1 Tg* worms are significantly enriched among genes downregulated following *KAT2* depletion in all three mammalian datasets (mouse, FDR = 4.82 × 10⁻⁵; keratinocytes, FDR = 3.39 × 10⁻⁵; MCF7Ras, FDR = 3.39 × 10⁻⁵) **(Fig. 2c–e)**. Conversely, orthogroups repressed by PCAF-1 are enriched among genes upregulated following KAT2 loss in mouse intestinal epithelial cells and human keratinocytes (FDR = 3.39 × 10⁻⁵ and 4.17 × 10⁻⁴, respectively), but this association is not significant in MCF7Ras cells (FDR = 0.50). Thus, our data indicate that genes induced by PCAF-1 in worms are consistently KAT2-dependent in mammalian cells, supporting conservation of KAT2-mediated transcriptional activation, whereas the conservation of PCAF-1 repressed programs is more context-dependent.

We next examined the biological functions associated with the conserved transcriptional response. Rank-based Gene Ontology enrichment analysis of PCAF-1 activated orthogroups identifies lipid metabolic and phosphorus metabolic processes among the shared biological processes, together with hydrolase activity and metal-ion binding among the shared molecular functions across all three mammalian datasets **(Fig. 2f–h)**. Given that protein kinase activity is enriched among PCAF-1-induced genes in worms **(Extended Data Fig. 1b)**, we asked whether kinase-encoding orthogroups also contribute to the conserved phosphorus-metabolism signal. Indeed, orthogroups corresponding to PCAF-1-induced protein kinases account for approximately 21–23% of the phosphorus-metabolism leading edge in each mammalian dataset, with several recurring across two or more datasets **(Extended Data Fig. 5)**. Beyond this shared core, additional enriched processes vary by cellular context, including xenobiotic responses in mouse intestinal epithelial cells, differentiation and small GTPase-mediated signaling in keratinocytes, and stress responses in MCF7Ras cells **(Fig. 2f–h)**. Several context-specific categories are also related to protein quality control, including proteolysis in keratinocytes and regulation of autophagy and ubiquitin protein ligase binding in MCF7Ras cells **(Fig. 2g,h)**. For example, the *hsp-3/HSPA5* orthogroup contributes to the MCF7Ras leading edge: *C. elegans hsp-3* is induced in *pcaf-1 Tg* animals, whereas mammalian ortholog *HSPA5*, encoding the ER chaperone BiP, is reduced following KAT2B silencing. Together, these analyses indicate that KAT2 acetyltransferases regulate a conserved core of metabolic and kinase-associated genes across species, while engaging protein quality-control pathways in a context-dependent manner.

### PCAF-1 reinforces protein quality control under stress and during aging

The enrichment of proteolysis-, autophagy-, and ubiquitin-associated functions among mammalian KAT2-dependent genes **(Fig. 2f–h)**, together with the concordant regulation of the *hsp-3/HSPA5* orthogroup, raised the possibility that PCAF-1 regulates protein homeostasis. To test this in *C. elegans*, we examined PCAF-1-responsive genes across the proteostasis network, including the endoplasmic reticulum (ER) unfolded protein response, ER-associated degradation (ERAD), molecular chaperones, the ubiquitin–proteasome system, and autophagy–lysosomal pathways. Elevated PCAF-1 significantly induces 70 proteostasis-associated genes spanning all of these pathways and represses 14 genes encoding negative regulators of protein quality control, with consistent changes across the three biological replicates **(Fig. 3a)**. Thus, PCAF-1 broadly remodels the expression of proteostasis genes in a direction predicted to enhance protein quality control, prompting us to examine how PCAF-1 affects proteostasis under stress and during aging.

**Fig. 3.**
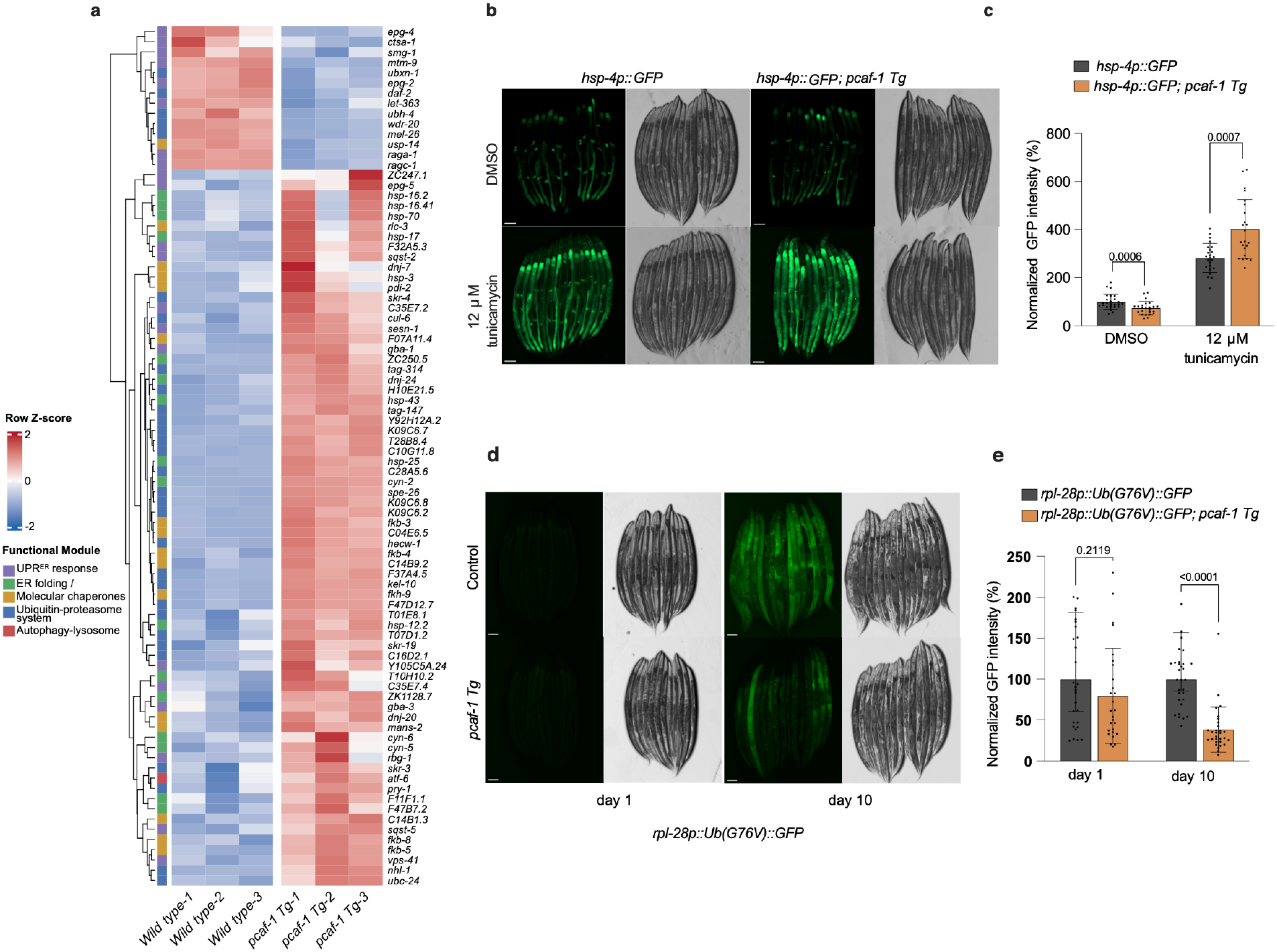
PCAF-1 reinforces protein quality control under stress and during aging. **a,** Targeted heatmap showing 70 significantly induced proteostasis-associated genes and 14 selected repressed genes annotated with negative regulatory functions in protein-quality-control pathways. Genes were identified from RNA-seq comparing wild-type and *pcaf-1 Tg* animals. Expression values are shown as row-wise Z-scores across three biological replicates per genotype. Functional annotations include ER unfolded protein response, ER folding/ER-associated degradation, molecular chaperones, ubiquitin–proteasome and autophagy–lysosomal pathways. **b,c,** Representative images (**b**) and quantification (**c**) of *hsp-4p::GFP* fluorescence in control and *pcaf-1 Tg* animals treated with DMSO or 10 µg ml⁻¹ tunicamycin (approximately 12 µM). GFP intensity was normalized to the DMSO-treated control. Data are from two independent biological experiments, with approximately 12 animals per condition per experiment. Bars show mean ± s.d., with individual animals shown as dots. Statistical significance was assessed using ordinary two-way ANOVA on log-transformed values followed by Šídák’s multiple-comparisons test. **d,e,** Representative images (**d**) and quantification (**e**) of *rpl-28p::Ub(G76V)::GFP* fluorescence in control and *pcaf-1 Tg* animals on days 1 and 10 of adulthood. Fluorescence was normalized to age-matched controls. Data are from two independent biological experiments, with approximately 12–15 animals per genotype and age per experiment. Bars show mean ± s.d. Statistical significance was assessed using ordinary two-way ANOVA followed by Šídák’s multiple-comparisons test. Scale bars, 100 µm (b,d).

The ER unfolded protein response (UPR^ER^) is of particular interest because mammalian KAT2B maintains UPR^ER^ gene expression and supports pancreatic β-cell adaptation to metabolic stress ^14^. We therefore asked whether PCAF-1 influences ER stress-responsive transcription using the *hsp-4p::GFP* reporter. *hsp-4* encodes an ER-resident chaperone homologous to mammalian BiP/GRP78, and its induction during ER stress depends on the IRE-1–XBP-1 pathway^25^. Under basal conditions, *pcaf-1 Tg* animals exhibit lower reporter fluorescence than controls. Upon exposure to 10 µM tunicamycin, an inhibitor of N-linked glycosylation that induces ER stress, reporter fluorescence increases approximately 2.8-fold in control animals but 5.5-fold in *pcaf-1 Tg* animals relative to their respective vehicle-treated levels **(Fig. 3b,c)**. Thus, increased PCAF-1 dosage lowers basal *hsp-4* expression while roughly doubling its stress-induced activation, consistent with reduced baseline ER stress and enhanced UPR^ER^ responsiveness.

We next asked whether PCAF-1 also influences the age-dependent accumulation of a proteasome substrate. For this, we used the ubiquitously expressed *rpl-28p::Ub(G76V)::GFP* reporter, in which a non-cleavable ubiquitin fusion targets GFP for ubiquitin-dependent proteasomal degradation^26,27^. Reporter fluorescence intensity is comparable between control and *pcaf-1 Tg* animals on day 1 of adulthood, but by day 10, *pcaf-1 Tg* animals exhibited ∼62% lower Ub(G76V)::GFP fluorescence than age-matched controls **(Fig. 3d,e)**. Conversely, *pcaf-1* RNAi increases reporter fluorescence by ∼37% on day 4 of adulthood, although this difference is no longer significant by day 13 **(Extended Data Fig. 6a,b)**. These findings indicate that the accumulation of a ubiquitin-proteasome substrate is sensitive to PCAF-1 dosage, with elevated PCAF-1 preventing its age-dependent build-up.

Together, these experiments show that elevated PCAF-1 enhances UPR^ER^ inducibility and limits age-associated accumulation of a proteasome substrate, supporting a role for PCAF-1 in maintaining protein homeostasis under stress and during aging. Because declining proteostasis capacity with age contributes to the accumulation of misfolded and aggregation-prone proteins^28,29^, we next investigated whether elevated PCAF-1 can protect against proteotoxicity in *C. elegans* models of human neurodegenerative diseases.

### PCAF-1 mitigates age-dependent protein aggregation and proteotoxicity

Given that PCAF-1 enhances UPR^ER^ inducibility and limits the age-associated accumulation of a proteasome substrate **(Fig. 3b-e)**, we tested whether it also protects against age-dependent protein aggregation. We first used animals expressing polyglutamine, which is a defining molecular feature of nine neurodegenerative disorders, including Huntington’s disease^30^. 35 glutamines (Q35) mark a critical threshold where proteins transition into an aggregation-prone state^28^. Thus, we used a Q35::YFP strain that expresses Q35 in body wall muscle under the *unc-54* promoter, which undergoes age-dependent aggregation, forming discrete fluorescent foci associated with progressive loss of motility^28^. *pcaf-1 Tg;Q35::YFP* animals do not show changes in aggregate number on day 1 of adulthood, but exhibit ∼36% and ∼30% significantly fewer Q35::YFP foci on days 4 and 7, respectively, compared with controls **(Fig. 4a,b).** To examine whether reducing endogenous *pcaf-1* produced the reciprocal phenotype, we measured aggregation following *pcaf-1 RNAi*. Conversely, *pcaf-1* knockdown does not affect aggregate numbers on day 1 but significantly increases Q35::YFP foci on days 4 and 7 **(Fig. 4c,d**). These opposing effects indicate that polyglutamine aggregate accumulation during adulthood is sensitive to PCAF-1 dosage. Moreover, reduced aggregation in *pcaf-1 Tg;Q35::YFP* animals is accompanied by significantly higher body-bend rates on days 1, 4, and 7 **(Fig. 4e)** and a ∼36% increase in median lifespan relative to Q35 controls **(Fig. 4f)**.

**Fig. 4.**
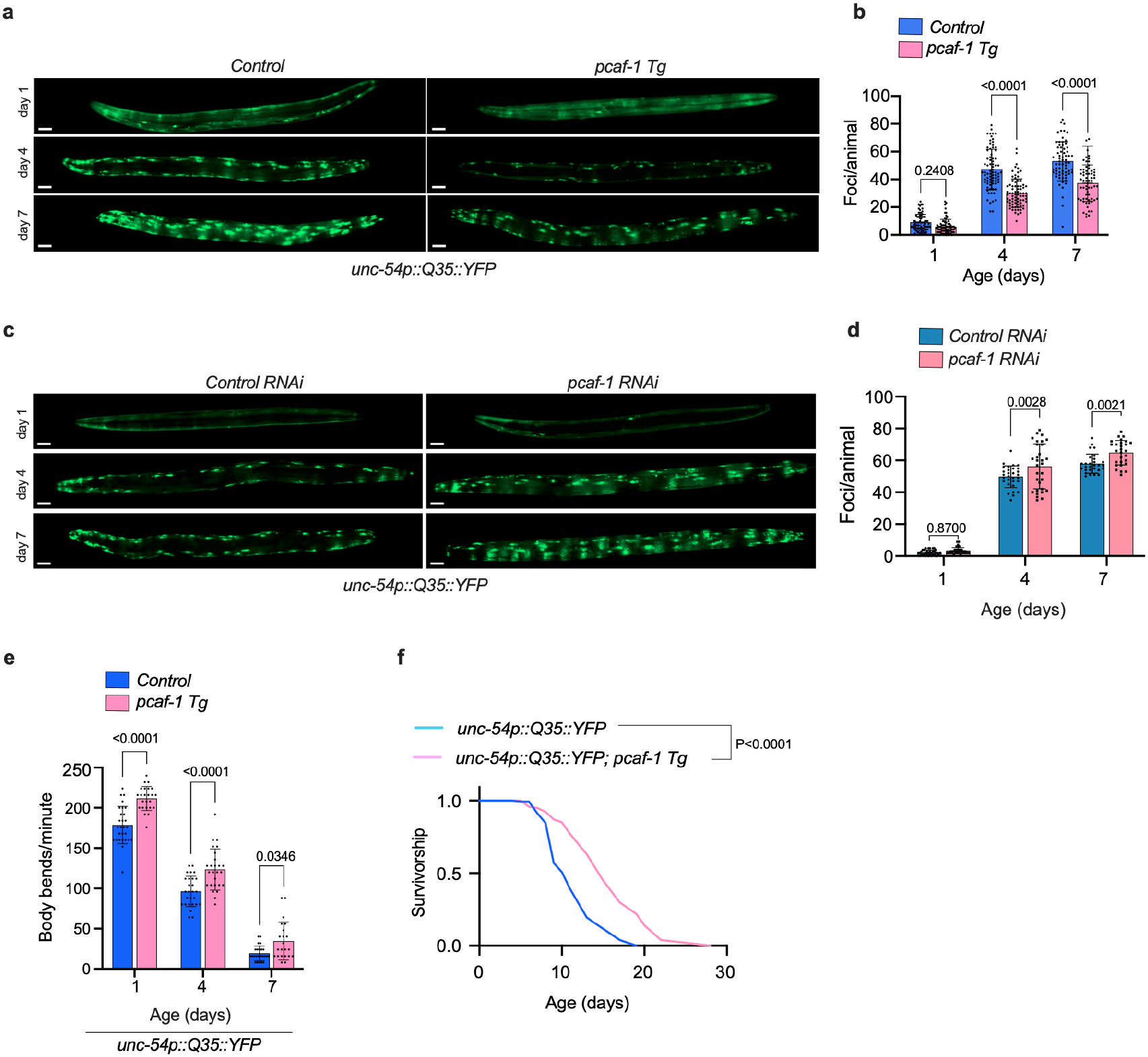
PCAF-1 suppresses polyglutamine aggregation and preserves motility and survival in a muscle proteotoxicity model. **a,b,** Representative fluorescence images (**a**) and quantification (**b**) of Q35::YFP aggregates in *C. elegans* expressing *unc-54p::Q35::YFP* (AM140) and animals carrying the *pcaf-1* transgene (Q35;*pcaf-1 Tg*) on days 1, 4 and 7 of adulthood. Aggregates were quantified as discrete fluorescent foci throughout the entire body of each animal. Approximately 15 animals per genotype at each age were examined in each of four independent biological experiments. Bars represent mean ± s.d., with individual animals shown as dots. Statistical significance was assessed using ordinary two-way ANOVA followed by Šídák’s multiple-comparisons test (day 1, adjusted *P* = 0.2408; days 4 and 7, adjusted *P* < 0.0001). **c,d,** Representative fluorescence images (**c**) and quantification (**d**) of Q35::YFP aggregates following control or *pcaf-1* RNAi on days 1, 4 and 7 of adulthood. Aggregates were quantified throughout the entire body of each animal. Approximately 15 animals per RNAi condition at each age were examined in each of two independent biological experiments. Bars represent mean ± s.d., with individual animals shown as dots. Statistical significance was assessed using ordinary two-way ANOVA followed by Šídák’s multiple-comparisons test (day 1, adjusted *P* = 0.8700; day 4, adjusted *P* = 0.0028; day 7, adjusted *P* = 0.0021). Scale bars, 50 µm (a,c). **e,** Locomotor activity of Q35 control and Q35;*pcaf-1 Tg* animals on days 1, 4 and 7 of adulthood, measured as body bends per minute. Approximately 14 animals per genotype at each age were examined in each of two independent biological experiments. Bars represent mean ± s.d., with individual animals shown as dots. Statistical significance was assessed using ordinary two-way ANOVA followed by Šídák’s multiple-comparisons test (days 1 and 4, adjusted *P* < 0.0001; day 7, adjusted *P* = 0.0346). **f,** Survival curves of Q35 control and Q35;*pcaf-1 Tg* animals. Transgenic animals exhibited a 36% increase in median lifespan. Survival curves were compared using the log-rank (Mantel–Cox) test (*P* < 0.0001).

To determine whether this protection extends beyond polyglutamine, we examined a structurally distinct aggregation-prone protein, human α-synuclein, whose aggregation is a pathological hallmark of Parkinson’s disease^31,32^. Animals expressing α-synuclein::YFP in body-wall muscle develop fluorescent inclusions during aging, accompanied by aggregation-associated functional decline^32,33^. As in the Q35 model, *pcaf-1 Tg* animals do not show changes in aggregate numbers on day 1 but reduce α-synuclein::YFP foci by ∼43% and ∼34% on days 4 and 7, respectively **(Fig. 5a,b)**. In contrast to Q35, however, *pcaf-1* RNAi does not significantly alter α-synuclein aggregation at any age examined **(Fig. 5c,d)**, suggesting that endogenous PCAF-1 is dispensable for limiting α-synuclein aggregation, or that its loss is compensated in this context. α-synuclein;*pcaf-1 Tg* animals also exhibit significantly higher body-bend rates on days 1, 4, and 7 of adulthood **(Fig. 5e**), and a ∼ 50% increase in median lifespan **(Fig. 5f)**. Together, our findings show that elevated PCAF-1 suppresses the age-dependent accumulation of two structurally distinct aggregation-prone proteins and preserves motility and survival in both models, indicating that PCAF-1 confers broad protection against age-associated proteotoxicity.

**Fig. 5.**
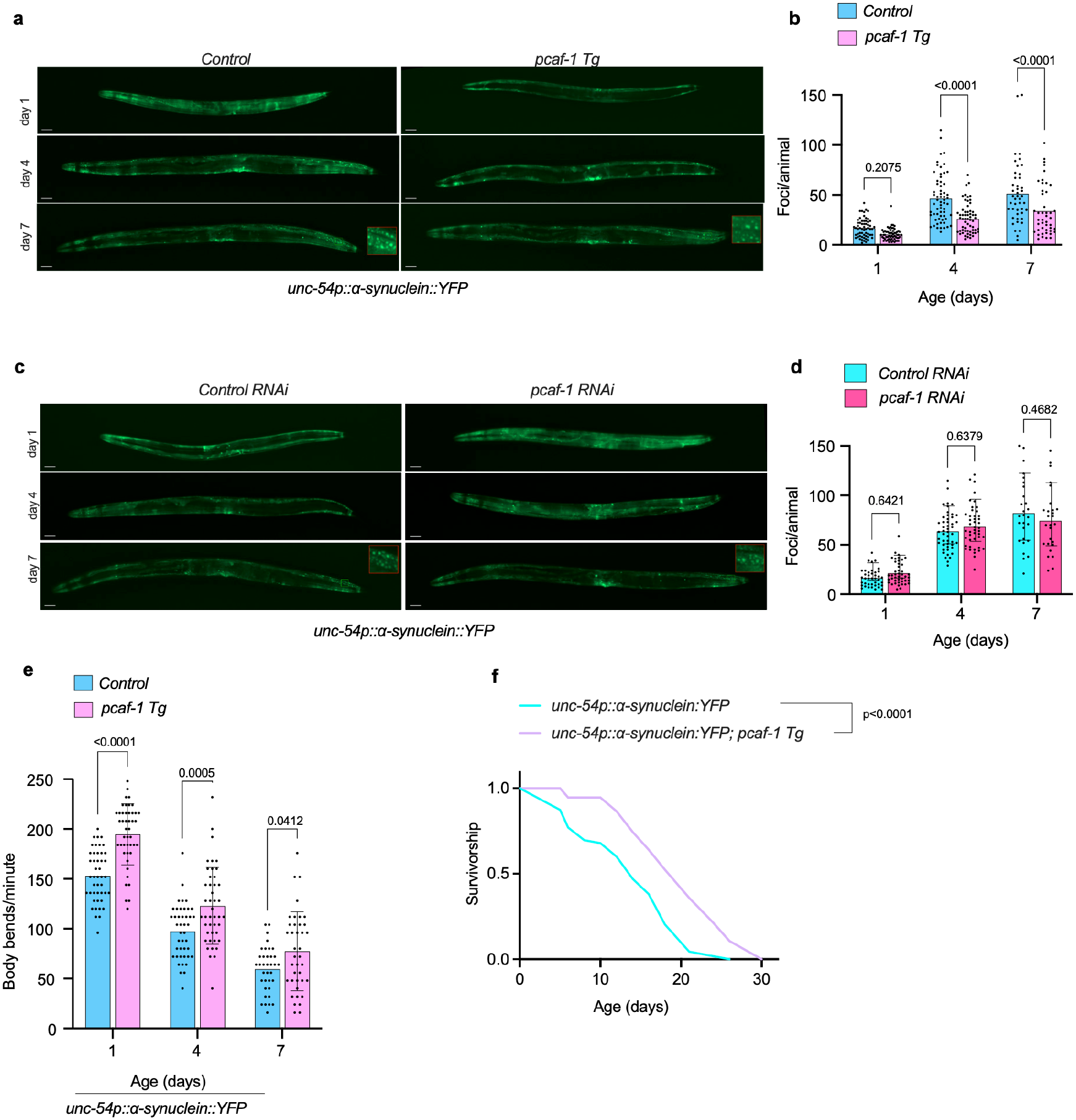
PCAF-1 protects against α-synuclein aggregation and extends healthspan in a Parkinson’s disease model. **a,b,** Representative fluorescence images (**a**) and quantification (**b**) of α-synuclein::YFP aggregates in *C. elegans* expressing *unc-54p::α-synuclein::YFP* (NL5901) and animals carrying the *pcaf-1* transgene (α-synuclein;*pcaf-1 Tg*) on days 1, 4 and 7 of adulthood. Aggregates were quantified as discrete fluorescent foci throughout the entire body of each animal. Approximately 15 animals per genotype at each age were examined in each of four independent biological experiments. Bars represent mean ± s.d., with individual animals shown as dots. Statistical significance was assessed using ordinary two-way ANOVA followed by Šídák’s multiple-comparisons test (day 1, adjusted *P* = 0.2075; days 4 and 7, adjusted *P* < 0.0001). **c,d,** Representative fluorescence images (**c**) and quantification (**d**) of α-synuclein::YFP aggregates following control or *pcaf-1* RNAi on days 1, 4 and 7 of adulthood. Aggregates were quantified throughout the entire body of each animal. Approximately 15 animals per RNAi condition at each age were examined in each of three independent biological experiments. Bars represent mean ± s.d., with individual animals shown as dots. Statistical significance was assessed using ordinary two-way ANOVA followed by Šídák’s multiple-comparisons test (day 1, adjusted *P* = 0.6421; day 4, adjusted *P* = 0.6379; day 7, adjusted *P* = 0.4682). Scale bars, 50 µm (a,c). **e,** Locomotor activity of α-synuclein control and α-synuclein;*pcaf-1 Tg* animals on days 1, 4 and 7 of adulthood, measured as body bends per minute. Approximately 15 animals per genotype at each age were examined in each of three independent biological experiments. Bars represent mean ± s.d., with individual animals shown as dots. Statistical significance was assessed using ordinary two-way ANOVA followed by Šídák’s multiple-comparisons test (day 1, adjusted *P* < 0.0001; day 4, adjusted *P* = 0.0005; day 7, adjusted *P* = 0.0412). **f,** Survival curves of α-synuclein control and α-synuclein;*pcaf-1 Tg* animals. Transgenic animals exhibited a 50% increase in median lifespan. Survival curves were compared using the log-rank (Mantel–Cox) test (*P* < 0.0001).

### PCAF-1 extends lifespan through DAF-16/FOXO and HLH-30/TFEB

We next sought to identify signaling pathways and transcriptional regulators underlying PCAF-1-dependent longevity. We analyzed genes upregulated and downregulated in *pcaf-1 Tg* animals separately using WormExp, a *C. elegans*-specific enrichment tool that compares gene lists with published transcriptional signatures^34^. We found that upregulated genes are significantly enriched for signatures associated with nutrient, longevity, and stress adaptation, including reduced insulin/IGF-1 signaling (IIS), AMPK-associated responses, and DAF-16/FOXO regulation **(Fig. 6a)**. Among transcription factor-associated signatures, PCAF-1-depenedent upregulated genes are enriched for DAF-16, PQM-1 and HLH-30 associated signatures, whereas downregulated genes are enriched for LIN-54, SKN-1 and BLMP-1 **(Extended Data Fig. 8a)**. Consistent with these nutrient-responsive longevity signatures, *pcaf-1 Tg* animals show significantly reduced expression of several IIS components, including *daf-2*, *age-1*, *pdk-1* and *akt-1*, as well as TOR-pathway genes, including *let-363/TOR*, *daf-15/Raptor* and *rsks-1/S6K* **(Fig. 6b)**. Genes involved in translation initiation, elongation, rRNA processing and ribosome biogenesis are also downregulated **(Extended Data Fig. 7e)**. Because reduced TOR–S6K signaling and inhibition of protein synthesis extend lifespan in *C. elegans*^16,17,35^, these changes are consistent with a shift away from growth-promoting programs.

**Fig. 6.**
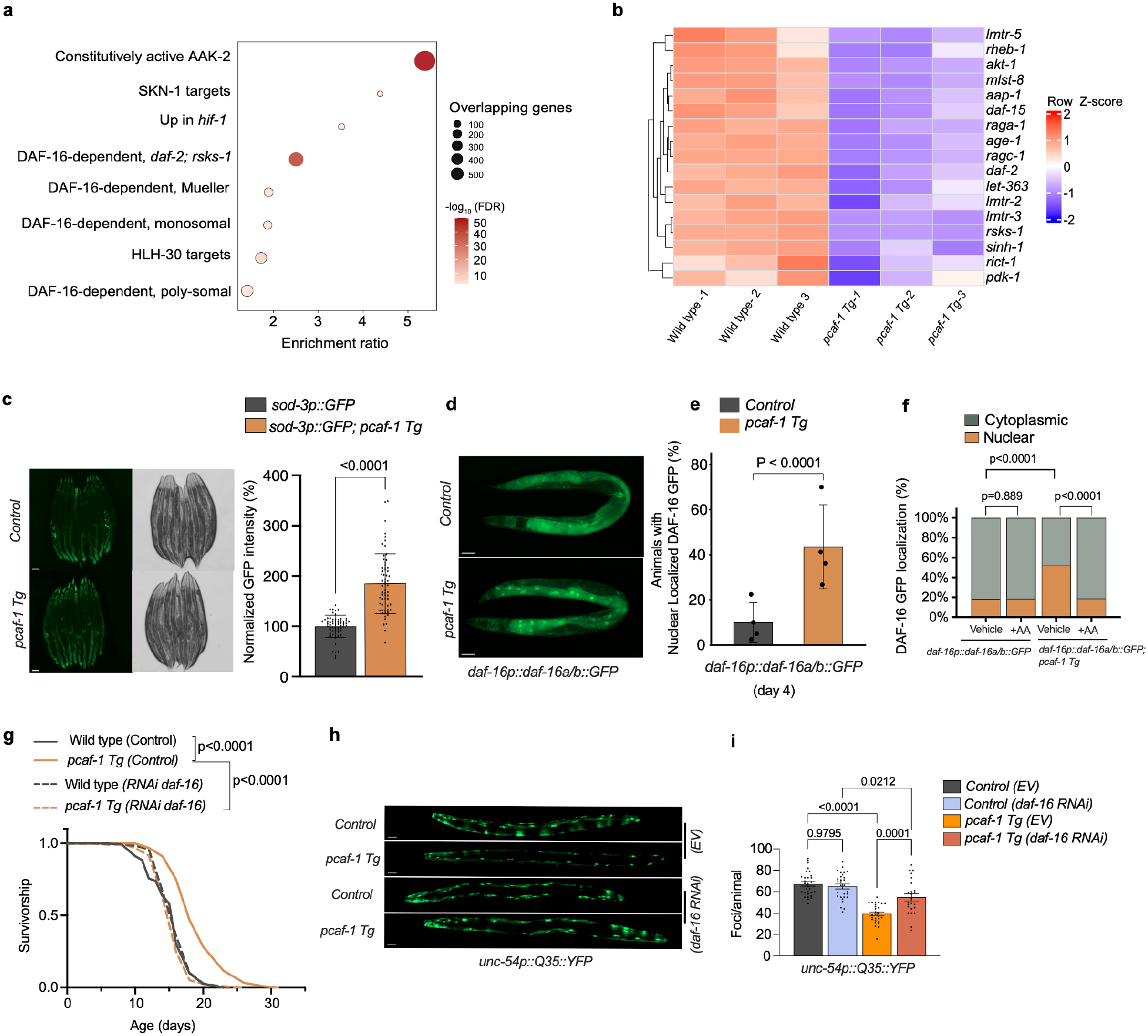
DAF-16/FOXO mediates PCAF-1-dependent longevity and proteostasis. **a,** WormExp enrichment of longevity- and stress-associated transcriptional signatures among genes upregulated in *pcaf-1 Tg* animals. Dot position represents the enrichment ratio, dot size indicates the number of overlapping genes, and dot color represents −log₁₀(FDR). Enrichment was assessed using the EASE-modified Fisher exact test with Benjamini–Hochberg correction. **b,** Heatmap of 17 significantly downregulated genes associated with insulin/IGF-1 signaling (IIS) and TOR signaling in wild-type and *pcaf-1 Tg* animals. Expression values are shown as row-wise Z-scores across three biological replicates per genotype. **c,** Representative fluorescence and bright-field images and quantification of *sod-3p::GFP* fluorescence in day-1 adults with or without *pcaf-1 Tg*. Approximately 20 animals per genotype per replicate were examined across three independent biological experiments. Bars show mean ± s.d., with individual animals shown as dots. Statistical significance was assessed using a two-sided Welch’s *t*-test, *P* < 0.0001. **d,e,** Representative fluorescence images (**d**) and quantification (**e**) of DAF-16::GFP nuclear localization in day-4 adults with or without *pcaf-1 Tg*. Animals were scored as exhibiting nuclear or cytoplasmic localization. Approximately 80 animals per genotype per replicate were examined across four independent biological experiments. Data are mean ± SD of replicate-level percentages. Cochran–Mantel–Haenszel (CMH) test, *P* = 4.72 × 10⁻²³. **f,** Nuclear and cytoplasmic localization of DAF-16::GFP in day-4 adults with or without *pcaf-1 Tg*, maintained on vehicle (ethanol) or 100 μM anacardic acid. Approximately 80 animals per condition per replicate were examined across two independent biological experiments with data pooled across the two experiments. Comparisons were performed using the CMH test (control vehicle versus control anacardic acid, *P* = 0.889; control vehicle versus *pcaf-1 Tg* vehicle, *P* < 0.0001; *pcaf-1 Tg* vehicle versus *pcaf-1 Tg* anacardic acid, *P* = 2.77 × 10⁻¹⁰). **g,** Lifespan of wild-type and *pcaf-1 Tg* animals maintained on control or *daf-16* RNAi from hatching. Representative Kaplan–Meier survival curves from two independent biological experiments (approximately 150 animals per condition per experiment). Survival was compared using the log-rank (Mantel–Cox) test. Wild-type with control RNAi versus *pcaf-1 Tg* with control RNAi, *P* < 0.0001; *pcaf-1 Tg* with control RNAi versus *pcaf-1 Tg* with *daf-16* RNAi, *P* < 0.0001. **h,i,** Representative fluorescence images (**h**) and quantification (**i**) of Q35::YFP aggregates in day-4 adults with or without *pcaf-1 Tg*, maintained on control EV or *daf-16* RNAi from hatching. Approximately 12–15 animals per condition were examined in each of two independent biological experiments. Bars show mean ± s.d., with individual animals shown as dots. Statistical significance was assessed using ordinary one-way ANOVA followed by Šídák’s multiple-comparisons test. Q35 control EV versus Q35 *daf-16* RNAi, adjusted *P* = 0.9795; Q35 control EV versus Q35;*pcaf-1 Tg* control EV, adjusted *P* < 0.0001; Q35;*pcaf-1 Tg* control EV versus Q35;*pcaf-1 Tg daf-16* RNAi, adjusted *P* = 0.0001; Q35 *daf-16* RNAi versus Q35;*pcaf-1 Tg daf-16* RNAi, adjusted *P* = 0.0212. Scale bars, 100 µm (c) and 50 µm (d,h).

In parallel, increased PCAF-1 dosage predominantly upregulated genes associated with reactive oxygen species metabolism, antioxidant activity, and oxidant detoxification **(Extended Data Fig. 8d)**. These include *sod-3*, which is induced 2.36-fold and encodes a mitochondrial manganese superoxide dismutase that is a direct transcriptional target of DAF-16/FOXO^36^. Consistently, *sod-3p::GFP* expression is ∼85% higher in *pcaf-1 Tg* adults **(Fig. 6c)**, whereas *pcaf-1 RNAi* significantly reduces its expression **(Extended Data Fig. 8e, f)**. Similarly, *gst-4*, which encodes a glutathione S-transferase involved in cellular detoxification and oxidative-stress defense^37^, was induced 2.78-fold and *gst-4p::GFP* expression was ∼51% higher in *pcaf-1 Tg* adults **(Extended Data Fig. 8 b,c)**.

The enrichment of DAF-16-associated transcriptional signatures, together with induction of *sod-3*, prompted us to examine DAF-16 subcellular localization. On day 4, nuclear DAF-16::GFP is significantly observed in 43.5% of *pcaf-1 Tg* animals compared with 10.0% of controls **(Fig. 6d,e)**., and this difference persisted on day 7 (37.9% versus 5.5%; **Extended Data Fig. 7a)**. To test whether PCAF-1 acetyltransferase activity contributed to this phenotype, we treated animals with anacardic acid, a small-molecule inhibitor previously shown to inhibit mammalian PCAF/KAT2B activity^38^. Anacardic acid (100 μM) significantly reduces DAF-16::GFP nuclear localization in *pcaf-1 Tg* animals, largely eliminating the difference from controls observed under vehicle conditions **(Fig. 6f)**, suggesting that enhanced DAF-16::GFP nuclear localization depends on PCAF-1 acetyltransferase activity.

To determine whether DAF-16 was functionally required for PCAF-1-dependent longevity, we examined survival following *daf-16* RNAi. *daf-16* knockdown completely abolishes the lifespan extension observed in *pcaf-1 Tg* animals, reducing survival of these transgenic animals to control levels **(Fig. 6g)**. We next examined whether DAF-16 also contributes to PCAF-1-mediated proteostasis in the Q35::YFP model. *daf-16 RNAi* increases aggregate accumulation in *pcaf-1 Tg;Q35::YFP* animals, partially reversing their reduced aggregation **(Fig. 6h,i)**. Taken together, our findings identify DAF-16/FOXO as a key mediator of PCAF-1 function that is required for both PCAF-1-dependent lifespan extension and contributes to protection against polyglutamine aggregation.

Given the reduced expression of TOR-pathway genes in *pcaf-1 Tg* animals **(Fig. 6b)**, we next examined HLH-30/TFEB, which promotes autophagy and lifespan extension and translocates to the nucleus upon reduced TOR signaling^39–41^. Elevated PCAF-1 increases the proportion of animals with nuclear HLH-30::GFP, primarily by shifting animals into the intermediate localization category **(Extended Data Fig. 7b,c)**. Furthermore, *hlh-30* RNAi fully suppresses the lifespan extension in *pcaf-1 Tg* animals **(Extended Data Fig. 7 d)**, revealing HLH-30 as a second essential effector of PCAF-1-mediated longevity. DAF-16/FOXO and HLH-30/TFEB are known to act together as combinatorial transcription factors that promote stress resistance and longevity^42^ (Lin et al., 2018). The requirement for both factors therefore suggests that PCAF-1 engages this pro-longevity module.

Finally, our gene set enrichment analysis shows that genes upregulated in *pcaf-1 Tg* animals overlap most strongly with those induced by constitutively active AAK-2/AMPK (**Fig. 6a**)^43–45^, prompting us to examine the role of AMPK. In human cells, AMPK activation stimulates KAT2B, leading to KAT2B-dependent hyperacetylation that inhibits cervical cancer growth^13^, and in yeast, the AMPK ortholog Snf1 regulates Gcn5 by phosphorylation and transcriptional activity^46^. We therefore hypothesized that AAK-2/AMPK acts upstream of PCAF-1. We reasoned that if this were the case, *pcaf-1* depletion should suppress the lifespan extension observed in animals that overexpress AAK-2(1–321), a truncated, constitutively active form of the AMPK α2 catalytic subunit^43–45^. Surprisingly, *pcaf-1* RNAi does not alter the survival of these animals (**Extended Data Fig. 9a**). In contrast, *aak-2* deletion reduces the median lifespan of *pcaf-1 Tg* animals (**Extended Data Fig. 9b**), consistent with AAK-2 acting downstream of PCAF-1. In line with this, phosphorylated AAK-2 levels are moderately but significantly higher in *pcaf-1 Tg* animals across three independent experiments, with a mean increase of 11.2% **(Extended Data Fig. 9c,d).** Because protein kinase activity is enriched among PCAF-1-induced genes **(Extended Data Fig. 1b)**, we examined potential regulators of AAK-2/AMPK signaling in our RNA-seq dataset. Among these, the gene encoding the creatine-kinase-like effector ARGK-1, *argk-1*, shows an approximately 11-fold increase in transcript abundance in *pcaf-1 Tg* animals (**Extended Data Fig. 9e**). ARGK-1 promotes AAK-2 activation and is required for the longevity of *rsks-1/S6K* mutants^47^, and *rsks-1* expression is itself reduced in *pcaf-1 Tg* animals (**Fig. 6b**). Consistent with the RNA-seq data, *argk-1p::GFP* fluorescence was increased by approximately 84% in *pcaf-1 Tg* animals **(Extended Data Fig. 9f, g)**. Together, our data reverse the expected hierarchy. Rather than acting upstream of PCAF-1, AAK-2/AMPK functions downstream of it to promote longevity, potentially through ARGK-1-mediated AMPK activation. Collectively, our results identify PCAF-1 as a pro-longevity acetyltransferase that couples stress-responsive transcription to proteostasis, extending lifespan and protecting against proteotoxicity through a conserved program that requires AMPK, DAF-16/FOXO, and HLH-30/TFEB.

## DISCUSSION

Here, we identify PCAF-1, the sole KAT2-family acetyltransferase in *C. elegans*, as a regulator of stress-responsive transcription and organismal resilience during aging. Increased PCAF-1 dosage extends lifespan, preserves locomotor function, and protects against age-associated proteotoxicity, with DAF-16/FOXO and HLH-30/TFEB required for the longevity response. These findings extend the physiological roles of KAT2 acetyltransferases beyond transcriptional regulation and chromatin modification^11^, establishing them as positive regulators of longevity and proteostasis.

KAT2 acetyltransferases are deeply conserved, but their contribution to organismal aging has been examined almost exclusively in yeast. There, Gcn5 influences both replicative and chronological lifespan through effects on the mitochondrial retrograde response, genome stability, and starvation-induced stress programs^10,48–50^. Particularly, reduced KAT2 activity is often beneficial in these contexts. Gcn5 inhibition extends lifespan in yeast and human cells^10^, while in *Drosophila*, loss of dGcn5 enhances Tau aggregate clearance and ameliorates neurodegeneration by relieving GCN5-mediated inhibition of the autophagy regulator TFEB^9^. In mammals, KAT2 function has been studied mainly in metabolic, stress adaptation, cancer, and cardiovascular diseases rather than aging. For instance, KAT2B sustains unfolded protein response gene expression as pancreatic β-cells adapt to metabolic challenge^14^, leaving the role of KAT2 acetyltransferases in metazoan aging largely undefined. Against this background, our finding that elevating PCAF-1 is protective is unexpected. Nonetheless, our cross-species transcriptomic analysis shows that the gene programs induced by PCAF-1 in worms are conserved targets of mammalian KAT2 regulation, despite the expansion from a single worm PCAF-1 protein to two mammalian paralogs. This opposing directionality, beneficial GCN5 loss in yeast and flies versus beneficial PCAF-1 gain in worms, may reflect substrate- and context-specific wiring: whereas GCN5 inhibits TFEB^9^, PCAF-1-mediated longevity requires HLH-30/TFEB and DAF-16/FOXO, suggesting that KAT2 enzymes can engage the autophagy–lysosomal network in opposite directions across species.

A central consequence of increased PCAF-1 dosage is protection against proteotoxicity during aging, when declining protein quality control allows misfolded and aggregation-prone proteins to accumulate^29,51^. In *pcaf-1 Tg* animals, reduced polyglutamine and α-synuclein aggregation is accompanied by preserved locomotor function and extended survival, indicating that protection is not specific to a single substrate. PCAF-1 also lowers the age-dependent accumulation of a proteasome-targeted reporter and heightens the inducibility of an ER stress-response reporter, pointing to effects across distinct arms of the proteostasis network. This breadth parallels the requirement for mammalian KAT2B in sustaining unfolded protein response gene expression during pancreatic β-cell adaptation to metabolic stress^14^.

DAF-16/FOXO provides a central link between PCAF-1, longevity, and proteostasis. DAF-16 coordinates transcriptional programs for metabolism, cellular defense, and protein homeostasis^52–56^, and in *pcaf-1 Tg* animals it shows sustained nuclear localization during adulthood, accompanied by induction of its direct target *sod-3*, consistent with enhanced DAF-16-dependent transcription. That *daf-16* knockdown abolishes the lifespan extension and partially reverses the polyglutamine protection establishes DAF-16 as functionally required for PCAF-1-dependent longevity and as a contributor to its protection against polyglutamine aggregation. Mammalian KAT2B/PCAF interacts with FOXO1 and promotes its nuclear accumulation, providing a precedent for the KAT2B-FOXO relationship^57^. Supporting this, anacardic acid substantially reduced PCAF-1-associated DAF-16 nuclear localization, suggesting that acetyltransferase-sensitive regulation contributes to this phenotype.

Acetylation is itself a key regulatory input for FOXO-family factors, complicating a simple model in which PCAF-1 directly activates DAF-16. In mammals, CBP/p300 and KAT2B/PCAF acetylate FOXO proteins, while SIRT1 deacetylates them and, in most contexts, acetylation attenuates FOXO activity by reducing DNA binding and transactivation, whereas SIRT1-mediated deacetylation restores activity^58–61^. In *C. elegans*, CBP-1 (KAT3 family) acetylates DAF-16; hence, *cbp-1* inhibition yields hypoacetylated, constitutively nuclear DAF-16^62^. Based on these works, *C. elegans* PCAF-1 may act indirectly by acetylating substrates upstream of DAF-16. However, because we did not directly measure DAF-16 acetylation, distinguishing these possibilities will require targeted analysis of DAF-16 acetylation in future work.

The requirement for AAK-2/AMPK places PCAF-1 within a canonical energy-sensing longevity network, but in an unexpected position. In *C. elegans*, AMPK is a classical upstream driver of lifespan^43–45^, and in other systems it acts upstream of KAT2 enzymes: human AMPK stimulates KAT2B, while yeast Snf1 regulates Gcn5^12,13,46^. Our epistasis data show the reverse relationship: *aak-2* loss shortens the lifespan of *pcaf-1 Tg* animals, whereas *pcaf-1* depletion does not affect animals with constitutively active AAK-2^44,45^, placing AMPK downstream of PCAF-1. One likely route is transcriptional. PCAF-1 strongly induces *argk-1*, a creatine-kinase-like effector that activates AAK-2 and is required for rsks-1/S6K-mutant longevity^47^, and *rsks-1* is also reduced in *pcaf-1 Tg* animals. Because the increase in phosphorylated AAK-2 is significant but modest and the link to ARGK-1 is correlative, future experiments will focus on the direction of regulation between PCAF-1 and AMPK.

In summary, our study links increased PCAF-1 dosage to stress-responsive transcription, reduced protein aggregation, and preserved organismal function, identifying KAT2-family acetyltransferases as positive regulators of longevity. Their conserved transcriptional signature suggests relevance beyond *C. elegans*. Our findings reframe the KAT2 family, previously characterized mainly as a negative regulator of autophagy^9^ and as a target whose inhibition extends lifespan in other systems^10^, as a dosage-sensitive promoter of proteostasis and healthy aging. Whether enhancing KAT2 activity confers similar benefits in mammalian tissues, and whether this can be achieved pharmacologically, will be important questions for translating these findings toward combating age-related proteotoxic disease and promoting healthy aging.

## MATERIALS AND METHODS

### *C. elegans* strains, transgene generation and maintenance

*Caenorhabditis elegans* strains were maintained at 20 °C on standard nematode growth medium seeded with *Escherichia coli* OP50-1. N2 was used as the wild-type strain. Two independent *pcaf-1*-transgenic lines were generated by microinjecting wild-type animals with 20 ng µl⁻¹ pCS3 (*eft-3p::pcaf-1 cDNA::unc-54 3′ UTR*) and 50 ng µl⁻¹ pCFJ104 (*myo-3p::mCherry::unc-54 3′ UTR*). *pcaf-1 Tg* line 1 and *pcaf-1 Tg* line 2 carry the extrachromosomal arrays *gsgEx3* and *gsgEx1*, respectively. Animals carrying the arrays were identified by mCherry expression in the body-wall muscles. Unless otherwise indicated, *pcaf-1 Tg* refers to line 1. Reporter and proteotoxicity strains were crossed with *pcaf-1 Tg* to generate the strains used for the indicated assays.

Unless otherwise stated, synchronized populations were generated by allowing gravid adults to lay eggs for 2 h. L4 larvae were transferred to the appropriate experimental plates, and the following day was designated day 1 of adulthood. Complete genotypes and sources of all strains, including the reporter and proteotoxicity crosses, are provided in **Supplementary Table 1**.

### RNA interference

RNAi clones targeting *pcaf-1*, *daf-16*, *hlh-30* and *aak-2* were obtained from the Ahringer RNAi library. Plasmids were isolated, confirmed by Sanger sequencing and transformed into *E. coli* OP50(xu363). OP50(xu363) carrying the empty L4440 plasmid was used as the empty-vector (EV) control. Bacterial cultures were grown overnight at 37 °C in LB containing 100 µg ml⁻¹ carbenicillin, pelleted and concentrated fivefold in fresh LB without antibiotic. 60-mm NGM plates without antibiotic were seeded with 100 µl of concentrated culture and incubated at room temperature for 3 d. Bacterial lawns were subsequently treated with 100 µl of 0.1 M IPTG containing 100 µg ml⁻¹ carbenicillin and incubated overnight. Gravid adults were allowed to lay eggs directly on induced RNAi plates for 2 h and were then removed.

### Lifespan analysis

Lifespan experiments were performed at 20 °C. On day 1 of adulthood, approximately 150 animals per condition were transferred to six 60-mm plates, with 25 animals per plate. Animals were maintained on OP50-1 or the indicated RNAi treatment and transferred to fresh plates carrying the corresponding bacterial treatment every other day beginning on day 1. Survival was scored every other day. Animals that failed to respond to three gentle taps to the head and tail were scored as dead. Animals were censored because of plate contamination, leaving the NGM, internal hatching or loss of vulval integrity. Kaplan–Meier survival curves were generated in GraphPad Prism and compared using the log-rank (Mantel–Cox) test. Independent lifespan experiments were analyzed separately and were not pooled. Exact animal numbers and the number of independent experiments are reported in the corresponding figure legends and source data.

### Heat-stress survival

Synchronized day-1 adult wild-type and *pcaf-1 Tg* line 1 animals were placed on 60-mm NGM plates seeded with OP50-1 and maintained at 35 °C. Approximately 60 animals per genotype were examined in each experiment across at least three plates. Survival was assessed hourly by scoring the same animals throughout the assay. Animals that failed to respond to gentle prodding with a platinum-wire pick were scored as dead. Two independent biological experiments were performed.

### Locomotor activity

Locomotor activity was assessed as described previously with minor modifications^63^. Individual animals were transferred from culture plates into 50 µl of M9 buffer in a 96-well plate. Bends toward one side of the body were counted for 15 s. Counts were multiplied by four to obtain the rate per minute and by two to account for bending in both directions. Separate animals were examined at each time point.

### RNA-seq

Animals were synchronized by bleaching gravid adults. Approximately 400 young adult animals were used for each sample and transferred to fresh OP50-1-seeded plates. Animals were collected 13 h after L4 stage as day-1 adults, with three independent biological replicate samples collected for each genotype. Animals were collected in M9 buffer and washed twice with M9 to remove residual bacteria. After the final centrifugation, the supernatant was removed, QIAzol lysis reagent was added, and samples were stored at −80 °C. For RNA isolation, samples were thawed at 37 °C, glass beads were added, and the samples were vortexed. Samples were subsequently subjected to two cycles of snap-freezing in liquid nitrogen, thawing at 37 °C for 2 min, and vortexing. Total RNA was isolated using the miRNeasy Tissue/Cells Advanced Mini Kit (QIAGEN, 217604). RNA quality assessment, ribosomal-RNA depletion, library preparation, and sequencing were performed by Azenta Life Sciences. Libraries were sequenced on an Illumina platform using 150-base paired-end reads.

### Gene-expression and functional analysis

Sequence reads were trimmed using Trimmomatic v.0.36^64^ and aligned to the *Caenorhabditis elegans* WBcel235 reference genome using STAR v.2.5.2b^65^. Gene-level counts were generated using featureCounts from the Subread package v.1.5.2^66^, retaining uniquely mapped reads overlapping annotated exons. Differential expression between wild-type and *pcaf-1 Tg* animals was analyzed using DESeq2^67^. P values were calculated using the Wald test and adjusted for multiple testing using the Benjamini–Hochberg procedure. Principal-component analysis was performed using regularized-logarithm-transformed counts for the 500 genes with the highest variance across the six samples. Complete gene-level differential-expression results for the 14,822 genes tested are provided in **Supplementary Data 1**. Genes with an adjusted P < 0.05 and |log_2_ fold change| ≥ 1 were considered differentially expressed. For pathway and functional enrichment analyses, an extended differential-expression threshold of adjusted P < 0.05 and |log_2_ fold change| ≥ log_2_(1.2) was used. KEGG pathway^68^ and Gene Ontology^69^ biological-process and molecular-function over-representation analyses were performed separately for induced and repressed genes using one-sided hypergeometric tests. P values were adjusted using the Benjamini–Hochberg procedure, and an FDR < 0.05 was considered significant. Protein kinase-associated genes were identified using the Gene Ontology molecular-function term GO:0004672, and genes induced at adjusted P < 0.05 and log_2_ fold change ≥ 1 or repressed at adjusted P < 0.05 and log₂ fold change ≤ −1 were included in the targeted heatmaps. Proteostasis-associated genes were identified from annotations related to the ER unfolded-protein response, ER protein folding and ER-associated degradation, molecular chaperones, the ubiquitin–proteasome system and autophagy–lysosomal degradation. IIS–TOR and translation-associated gene sets were curated from established pathway components and Gene Ontology annotations. Heatmaps were generated from normalized expression values, standardized as row-wise Z-scores and clustered using Euclidean distance with complete linkage. Heatmaps were generated in R using ComplexHeatmap^70^ and circlize^71^. Oxidative-stress-associated genes were identified using Gene Ontology annotations related to reactive oxygen species metabolism, oxidant detoxification, antioxidant activity and responses to reactive oxygen species. The targeted volcano plot shows genes in this set with adjusted P < 0.05 and |log_2_ fold change| ≥ 1. The gene sets used for the kinase, proteostasis, IIS–TOR, translation and oxidative-stress analyses are provided in **Supplementary Data 1**.

### WormExp analysis

Enrichment for previously reported *C. elegans* transcriptional signatures was assessed using WormExp^34^. Genes induced in *pcaf-1 Tg* animals at adjusted P < 0.05 and log_2_ fold change ≥ 1 were analyzed using the 14,822 genes tested in the RNA-seq analysis as the custom background. Over-representation was assessed using the EASE-modified Fisher exact test implemented in WormExp, and terms with an FDR < 0.05 were considered significant. Transcription factor target gene sets were obtained from the WormExp TF Targets category. For this analysis, induced and repressed genes were analyzed separately using the extended differential-expression threshold defined above.

### Cross-species transcriptomic analysis

Publicly available transcriptomic datasets were obtained for primary mouse intestinal epithelial cells following combined *Kat2a* and *Kat2b* loss (GSE235789)^22^, human keratinocytes following *KAT2B* knockdown (GSE215244)^24^, and human MCF7Ras cells following *KAT2B* siRNA treatment (GSE195761)^23^. Genes were ranked according to their transcriptional response to KAT2 loss. The DESeq2 Wald statistic was used for the mouse and keratinocyte datasets, whereas MCF7Ras genes were ranked by signed −log₁₀(P value), with the sign determined by the direction of the log₂ fold change. Worm genes induced or repressed in *pcaf-1 Tg* animals at adjusted P < 0.05 and |log_2_ fold change| ≥ log_2_(1.2) were mapped to mouse or human orthologs using reciprocal-best orthology relationships from the Alliance of Genome Resource^72^. When multiple mammalian genes mapped to the same worm-anchored orthogroup, their ranking statistics were summarized using the median. The PCAF-1-induced and PCAF-1-repressed programs were tested against each ranked mammalian orthogroup dataset using the fgsea R/Bioconductor package with 100,000 permutations^73^. Benjamini–Hochberg correction was applied across the six program-by-dataset comparisons. For functional analysis of the conserved induced program, Gene Ontology biological-process and molecular-function subsets were tested against each complete orthogroup ranking using multilevel GSEA. P values were adjusted separately for each dataset and ontology using the Benjamini–Hochberg procedure, and an FDR < 0.05 was considered significant. For the phosphorus metabolic process (GO:0006793), leading-edge orthogroups were extracted separately from each mammalian dataset. Kinase-associated orthogroups were identified by overlap with the PCAF-1 induced protein kinase gene set defined above, and those contributing to the leading edge in at least two datasets were considered recurrent.

### Phylogenetic and protein-sequence analysis

Full-length amino-acid sequences of KAT2-family proteins were obtained from UniProtKB. For phylogenetic analysis, 22 sequences were aligned using MAFFT v.7.525 with the L-INS-i strategy^74^. Poorly aligned positions were removed using trimAl v.1.5.1 with the automated1 method^75^. Maximum-likelihood phylogenetic inference was performed using IQ-TREE v.2.4.0^76^. ModelFinder selected Q.insect+I+G4 as the best-fitting substitution model according to the Bayesian information criterion^77^. Branch support was assessed using 1,000 SH-like approximate-likelihood-ratio replicates and 1,000 ultrafast-bootstrap replicates with bootstrap nearest-neighbor-interchange optimization. The tree was rooted using *Saccharomyces cerevisiae* and *Arabidopsis thaliana* Gcn5 sequences. To compare sequence conservation among representative family members, seven full-length proteins were aligned using MAFFT v.7.526 with the E-INS-i strategy. Pairwise sequence identity was calculated using alignment positions containing amino acids in both sequences. Domain-specific comparisons used UniProtKB-annotated N-acetyltransferase and bromodomain boundaries. Protein lengths and domain annotations used for the domain-architecture comparison were obtained from UniProtKB.

### Fluorescence microscopy and image analysis

Fluorescence imaging was performed using a Zeiss Axio Zoom V16 fluorescence microscope equipped with an Axiocam 705 color camera and ZEN 3.6 Pro acquisition software. GFP and mCherry fluorescence images were acquired in monochrome mode. Animals were immobilized in 10 mM sodium azide as described previously with minor modifications^78^ and imaged directly on NGM plates. Images were acquired at ×50 magnification using the same acquisition settings for all groups compared within each biological experiment. Whole-animal mean GFP fluorescence was quantified using Fiji^79^. The outline of each animal was traced to define the region of interest, and corrected mean fluorescence was calculated by subtracting the mean fluorescence of a nearby background region from the mean whole-animal fluorescence. Values were normalized to the corresponding control group within each biological experiment. Separate animals were examined at each time point.

### Chemical treatments

Age-synchronized late-L4 *hsp-4p::GFP* animals were incubated for 2 h at room temperature in M9 containing 10 µg ml⁻¹ tunicamycin or DMSO vehicle, with gentle rotation, as described previously with minor modifications^78^. Animals were washed three times with M9, transferred to fresh OP50-1-seeded NGM plates and allowed to recover for approximately 16 h before imaging as day-1 adults. For anacardic acid treatment, DAF-16::GFP animals were exposed from hatching to OP50-1-seeded NGM plates containing 100 µM anacardic acid or ethanol vehicle. Treatment plates were prepared the day before use, and DAF-16::GFP localization was scored.

### Polyglutamine and α-synuclein aggregation assays

Q35::YFP aggregation was assessed in AM140 animals as described previously with minor modifications^80^. α-Synuclein::YFP aggregation was assessed in NL5901 animals^32^ using a whole-animal scoring approach^81^. Animals were imaged at ×50 magnification as described above. Aggregates were defined as discrete, bright fluorescent puncta distinguishable from the surrounding diffuse fluorescence and were counted manually throughout the entire body from saved fluorescence images. Separate animals were examined at each time point.

### DAF-16 and HLH-30 nuclear-localization assays

DAF-16::GFP and HLH-30::GFP localization was assessed in live, non-immobilized animals on food using a Zeiss Axio Zoom V16 fluorescence microscope at ×150 magnification. Because these transcription factors are sensitive to environmental stress and HLH-30::GFP can translocate to the nucleus in response to starvation or mounting stress^41^, animals were examined promptly in batches of approximately ten to minimize handling-induced changes in localization. DAF-16::GFP localization was classified as nuclear when any distinct nuclear GFP signal was observed and as cytosolic when no nuclear signal was detected. HLH-30::GFP localization was classified as nuclear when prominent nuclear GFP signal was observed throughout the intestine, intermediate when nuclear localization was restricted primarily to approximately four to five nuclei in the posterior intestine, or cytosolic when no distinct nuclear localization was detected. Where indicated, localization was assessed following anacardic-acid treatment as described above.

### Western blotting

Approximately 100 synchronized day-4 adults were washed twice with M9, resuspended in 2× Laemmli sample buffer and heated at 95 °C for 10 min. Whole-worm lysates were separated on 4–20% Mini-PROTEAN TGX precast gels (Bio-Rad, 4561093) and transferred to nitrocellulose membranes. Membranes were blocked overnight at 4 °C in 5% bovine serum albumin in TBST and probed with phospho-AMPKα (Thr172) antibody (Cell Signaling Technology, 2535T; 1:1,000 in 3% bovine serum albumin). After imaging, membranes were stripped for 10 min using Restore Western Blot Stripping Buffer (Thermo Scientific, 21059), washed three times with TBST, blocked for 1 h in 5% bovine serum albumin and reprobed with α-tubulin antibody (Abcam, ab52866; 1:1,000 in TBST). Primary and HRP-linked anti-rabbit IgG secondary-antibody incubations (Cell Signaling Technology, 7074; 1:200 in TBST) were performed for 1.5 h at room temperature. Signals were detected using SuperSignal West Pico PLUS Chemiluminescent Substrate (Thermo Fisher Scientific, 34577) and a ChemiDoc MP Imaging System (Bio-Rad). Band intensities were quantified using Fiji, and phospho-AAK-2 intensity was normalized to α-tubulin and expressed relative to the wild-type mean.

### Statistics and reproducibility

Fiji was used for image quantification, and GraphPad Prism was used for statistical analyses. Statistical details for individual experiments, including the statistical tests and representation of variability, are provided in the corresponding figure legends. Survival curves were compared using the log-rank (Mantel–Cox) test. Comparisons involving multiple groups or factors were performed using ordinary one-way or two-way analysis of variance followed by Šídák or Bonferroni multiple-comparisons tests, as indicated. Other comparisons were performed using the statistical tests specified in the corresponding figure legends. Differentially expressed genes were identified using the Wald test, as described above. For all experiments, *P* < 0.05 was considered statistically significant. No statistical method was used to predetermine sample sizes. No formal blinding was used. Lifespan and protein-aggregation assays were performed or scored by more than one investigator across independent experiments. No measurements were excluded, and lifespan censoring followed the criteria described above.

## Supporting information

Supplementary Table 1. C. elegans strains used in this study

Supplementary Data 1. Complete RNA-seq differential-expression results

Extended Data Figure 2. Increased PCAF-1 dosage alters the expression of protein kinase-associated genes

Extended Data Figure 4. Multiple-sequence alignment of GCN5/PCAF/KAT2-family proteins

## ACKNOWLEDGEMENTS

We thank all members of the Silva-García laboratory for their helpful discussions. Bristol N2 strain was provided by the Caenorhabditis Genetics Center (CGC), which is funded by the NIH Office of Research Infrastructure Programs (P40 OD010440).

## AUTHOR CONTRIBUTIONS

A.S. and C.G.S.G. designed the study. A.S. performed most experiments and analyzed results. K.N. performed the body-bend assays and western blot experiments. L.B.R. characterized the aggregation and survival phenotypes of the Q35 lines. J.T. assisted with image analysis and α-synuclein survival assays. E.D. generated the *pcaf-1* transgenic lines. M.H.T. assisted with RNA-seq sample collection. A.S. and C.G.S.G. wrote and revised the manuscript with input from all authors. All authors reviewed and approved the final manuscript.

---

**Extended Data Fig. 1.**
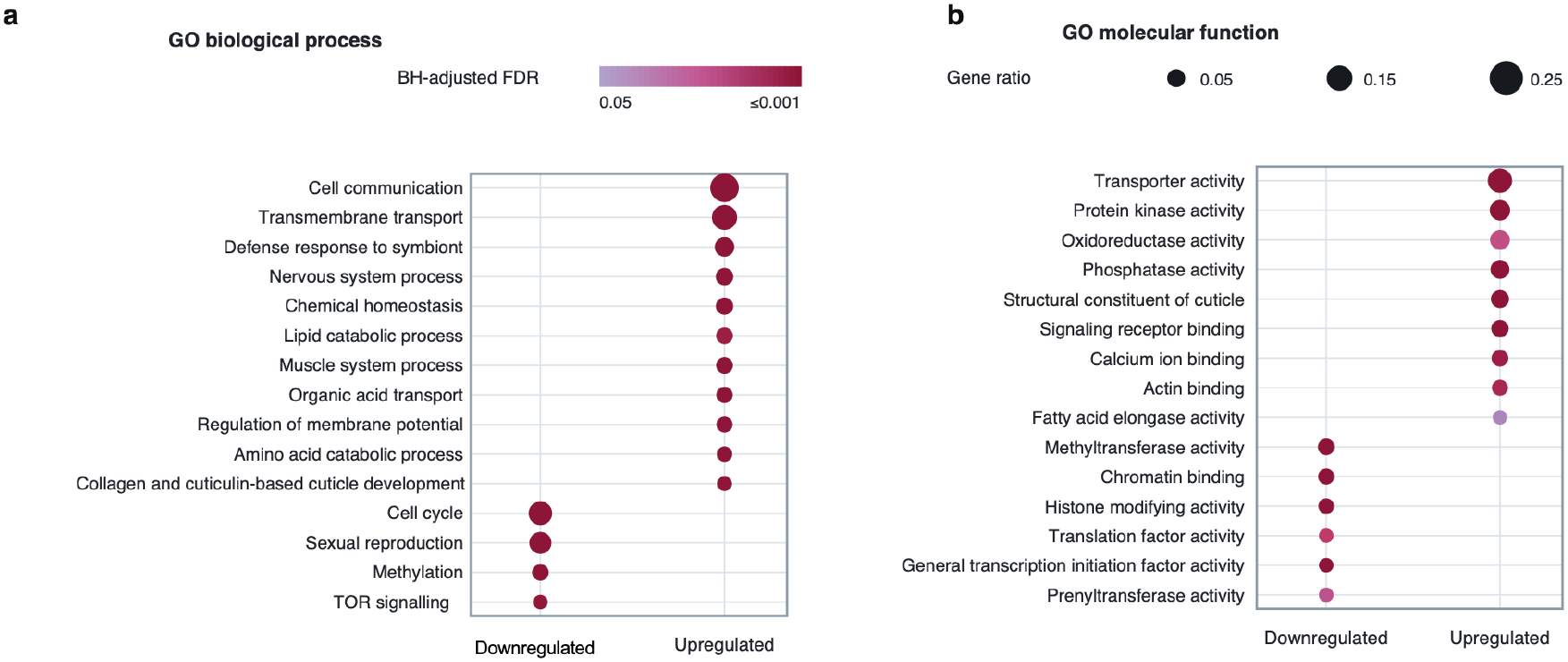
Gene Ontology analysis reveals PCAF-1-dependent changes in biological processes and molecular functions. **a,** Gene Ontology (GO) biological process enrichment analysis of genes upregulated and downregulated in *pcaf-1* transgenic (*pcaf-1 Tg*) animals relative to wild-type. PCAF-1-induced genes were enriched for processes associated with cell communication, transmembrane transport, defense responses, nervous and muscle system functions, chemical homeostasis, lipid catabolism, and regulation of membrane potential. Repressed genes were enriched for cell cycle, sexual reproduction, methylation, and TOR signaling. **b,** GO molecular function enrichment analysis of the same gene sets. Upregulated genes were enriched for transporter activity, protein kinase activity, oxidoreductase activity, phosphatase activity, and signaling receptor binding, whereas downregulated genes were enriched for methyltransferase activity, chromatin binding, histone-modifying activity, and translation factor activity. **a,b,** Enrichment analyses were performed separately for upregulated and downregulated genes using adjusted *P* < 0.05 and |log₂ fold change| ≥ 0.263 (equivalent to a 1.2-fold expression change). Selected significantly enriched GO (FDR < 0.05) terms are shown. Dot size represents the gene ratio, and dot color indicates the Benjamini–Hochberg-adjusted false discovery rate (FDR).

**Extended Data Fig. 2 is provided as a separate supplementary PDF.**

**Extended Data Fig. 2. Increased PCAF-1 dosage alters the expression of protein kinase-associated genes.**

**a,b.** Heatmaps showing the expression profiles of 127 significantly upregulated (**a**) and 13 significantly downregulated (**b**) protein kinase-associated genes in wild-type and *pcaf-1 Tg* animals. Protein kinase genes were identified using the Gene Ontology molecular function term GO:0004672 (protein kinase activity). Genes were selected based on an adjusted *P* < 0.05 and |log₂ fold change| ≥ 1 in *pcaf-1 Tg* relative to wild-type animals. RNA-seq data were obtained from three independent biological replicates per genotype. Expression values are displayed as row-wise Z-scores.

**Extended Data Fig. 3.**
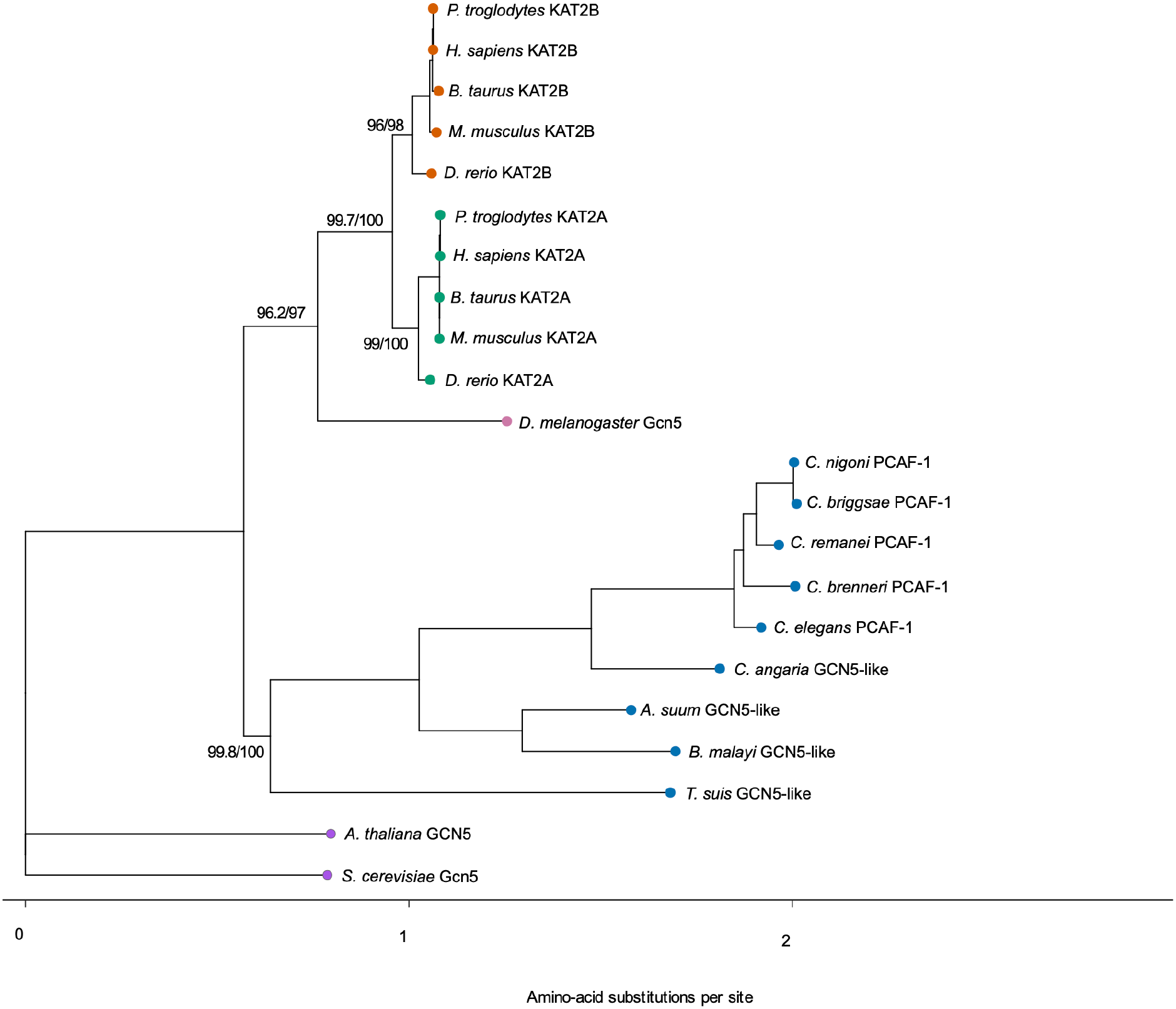
Evolutionary conservation of the GCN5/PCAF/KAT2 acetyltransferase family. Maximum-likelihood phylogenetic tree of 22 full-length GCN5/PCAF/KAT2-family protein sequences from nematodes, insects, vertebrates, plants and fungi. The analysis identifies *C. elegans* PCAF-1 within the conserved KAT2 family and distinguishes the vertebrate KAT2A and KAT2B paralog groups. Node labels indicate SH-aLRT/ultrafast-bootstrap support percentages based on 1,000 replicates each. Branch lengths represent the estimated number of amino-acid substitutions per site.

**Extended Data Fig. 4 is provided as a separate supplementary PDF.**

**Extended Data Fig. 4. Amino-acid sequence conservation among representative GCN5/PCAF/KAT2-family acetyltransferases.**

Multiple-sequence alignment of seven representative GCN5/PCAF/KAT2-family proteins from *Caenorhabditis elegans* (PCAF-1), *Saccharomyces cerevisiae* (Gcn5), *Drosophila melanogaster* (Gcn5), *Homo sapiens* (KAT2A and KAT2B), *Danio rerio* (KAT2B) and *Mus musculus* (KAT2B). Full-length protein sequences were aligned using MAFFT v7.526 with the E-INS-i strategy. The alignment reveals conserved amino acid residues, particularly within the catalytic acetyltransferase region. Within this region, *C. elegans* PCAF-1 shares 60.3% and 63.1% sequence identity with human KAT2A and KAT2B, respectively. Blue shading indicates positions containing identical residues across all seven proteins. Asterisks (*), colons (:) and periods (.) denote identical, strongly similar and weakly similar residues, respectively. Protein lengths and UniProt accession numbers are provided.

**Extended Data Fig. 5.**
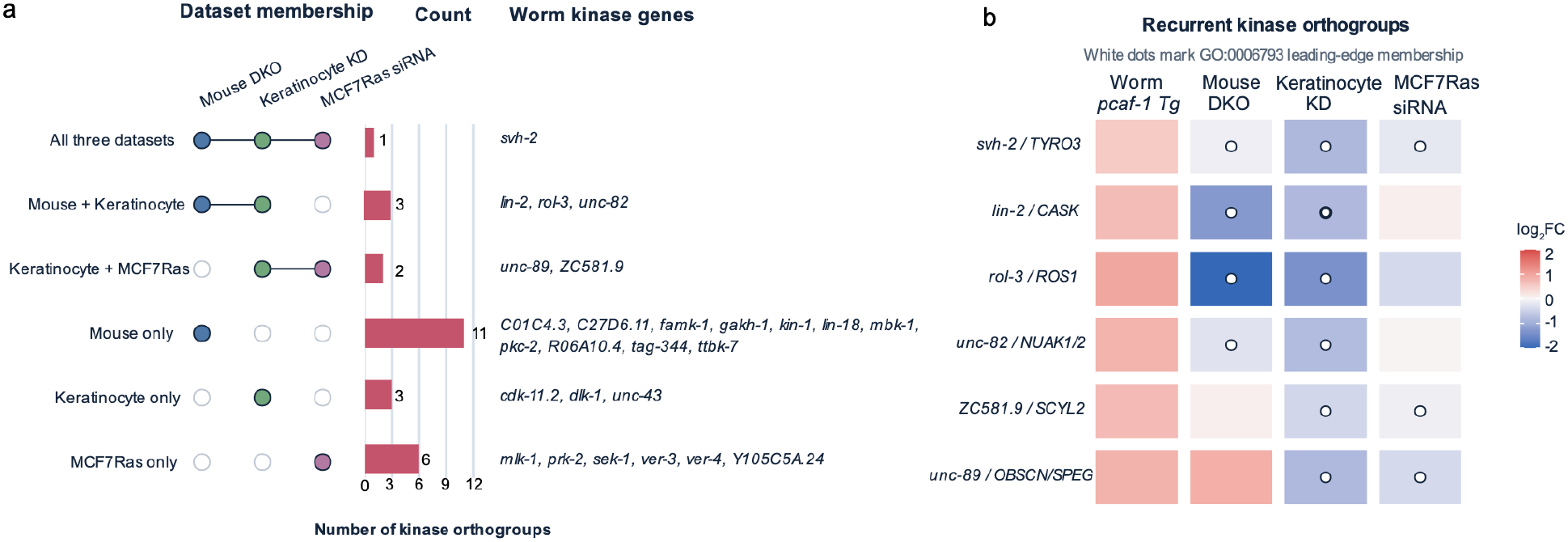
PCAF-1-induced kinase orthogroups contribute to conserved phosphorus-metabolism enrichment across mammalian KAT2-loss datasets. **a,** Dataset membership of PCAF-1-induced kinase-associated orthogroups contributing to the phosphorus metabolic process (GO:0006793) leading edge in mouse *Kat2a/Kat2b* double-knockout intestinal epithelial cells, human keratinocytes following KAT2B knockdown and human MCF7Ras cells following KAT2B siRNA treatment. Filled circles indicate the datasets in which each group of orthogroups contributes to the leading edge. Bars and adjacent numbers show the number of kinase orthogroups in each dataset intersection, and the corresponding *C. elegans* kinase genes are listed. **b**, Expression profiles of the six recurrent kinase orthogroups contributing to the leading edge in at least two mammalian datasets. Rows are labeled with the *C. elegans* gene and corresponding mammalian ortholog or orthologs. Color represents log₂ fold change in *pcaf-1 Tg* animals and the indicated mammalian KAT2-loss datasets, and white dots indicate leading-edge membership. PCAF-1-induced kinase-associated orthogroups accounted for approximately 21–23% of the phosphorus-metabolism leading edge in each mammalian dataset.

**Extended Data Fig. 6.**
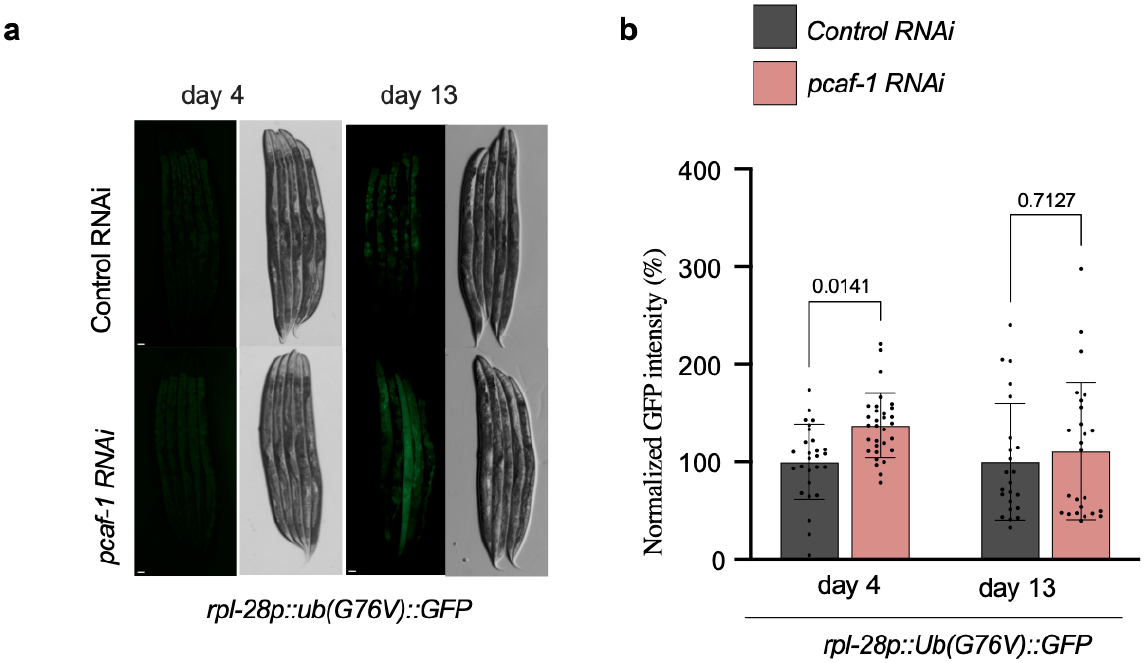
*pcaf-1* knockdown increases accumulation of a proteasome-targeted reporter during adulthood. **a,** Representative fluorescence and bright-field images of *rpl-28p::Ub(G76V)::GFP* animals treated with control or *pcaf-1* RNAi from hatching, imaged on days 4 and 13 of adulthood. **b,** Quantification of whole-worm mean GFP fluorescence normalized to age-matched control RNAi animals. Data are from two independent biological experiments, with approximately 12–15 animals per condition per experiment. Bars show mean ± s.d., with individual animals shown as dots. Statistical significance was assessed using ordinary two-way ANOVA followed by Šídák’s multiple-comparisons test. Scale bar, 50 µm.

**Extended Data Fig. 7.**
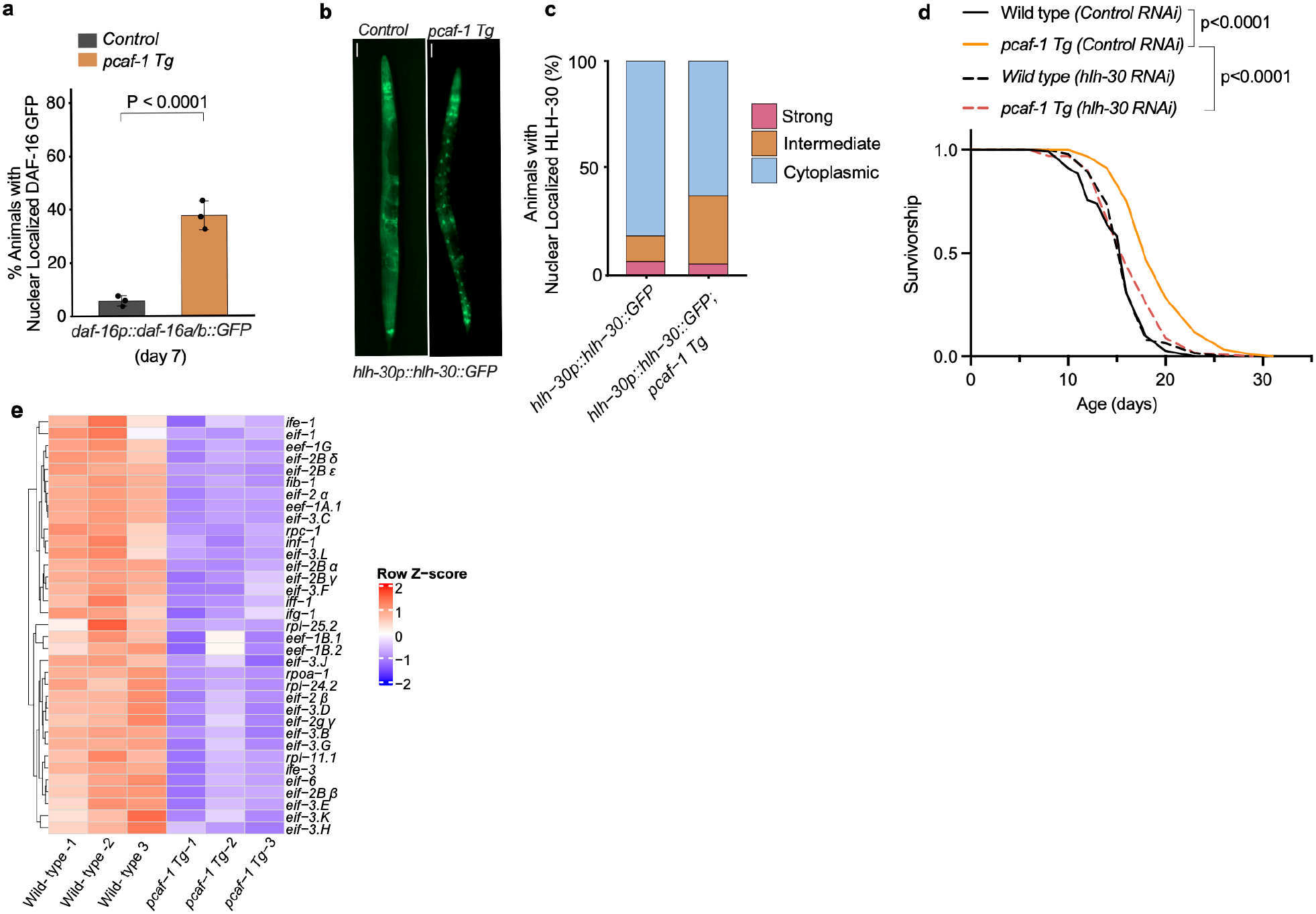
PCAF-1 promotes HLH-30 nuclear localization and reduces translation-associated gene expression. **a,** Nuclear localization of DAF-16::GFP in day-7 adults with or without *pcaf-1 Tg*. Approximately 40–70 animals per genotype were examined in each of three independent biological experiments. Day-4 and day-7 measurements were obtained from separate animals within the same synchronized experimental cohorts. Data are mean ± s.d. of replicate-level percentages. Statistical significance was assessed using the Cochran–Mantel–Haenszel (CMH) test (*P* = 1.20 × 10⁻¹¹). **b,c,** Representative fluorescence images (**b**) and quantification (**c**) of HLH-30::GFP localization in day-1 adults with or without *pcaf-1 Tg*. Animals were categorized as having cytoplasmic, intermediate nuclear or strongly nuclear localization. Approximately 80 animals per genotype were examined in each of six independent biological experiments. Bars show the percentage of animals in each localization category. The combined strong and intermediate nuclear categories were compared with cytoplasmic localization using the CMH test (*P* = 1.25 × 10⁻¹⁰). **d,** Lifespan of wild-type and *pcaf-1 Tg* animals maintained on control or *hlh-30* RNAi from hatching. A representative experiment from two independent biological experiments is shown, with approximately 150 animals per condition in each experiment. Survival was compared using the log-rank (Mantel–Cox) test (wild type with control RNAi versus *pcaf-1 Tg* with control RNAi, *P* < 0.0001; *pcaf-1 Tg* with control RNAi versus *pcaf-1 Tg* with *hlh-30* RNAi, *P* < 0.0001). **e,** Heatmap of 35 significantly downregulated genes associated with translation initiation, translation elongation, rRNA processing and ribosome biogenesis in wild-type and *pcaf-1 Tg* animals. Expression values are shown as row-wise Z-scores across three biological replicates per genotype.

**Extended Data Fig. 8.**
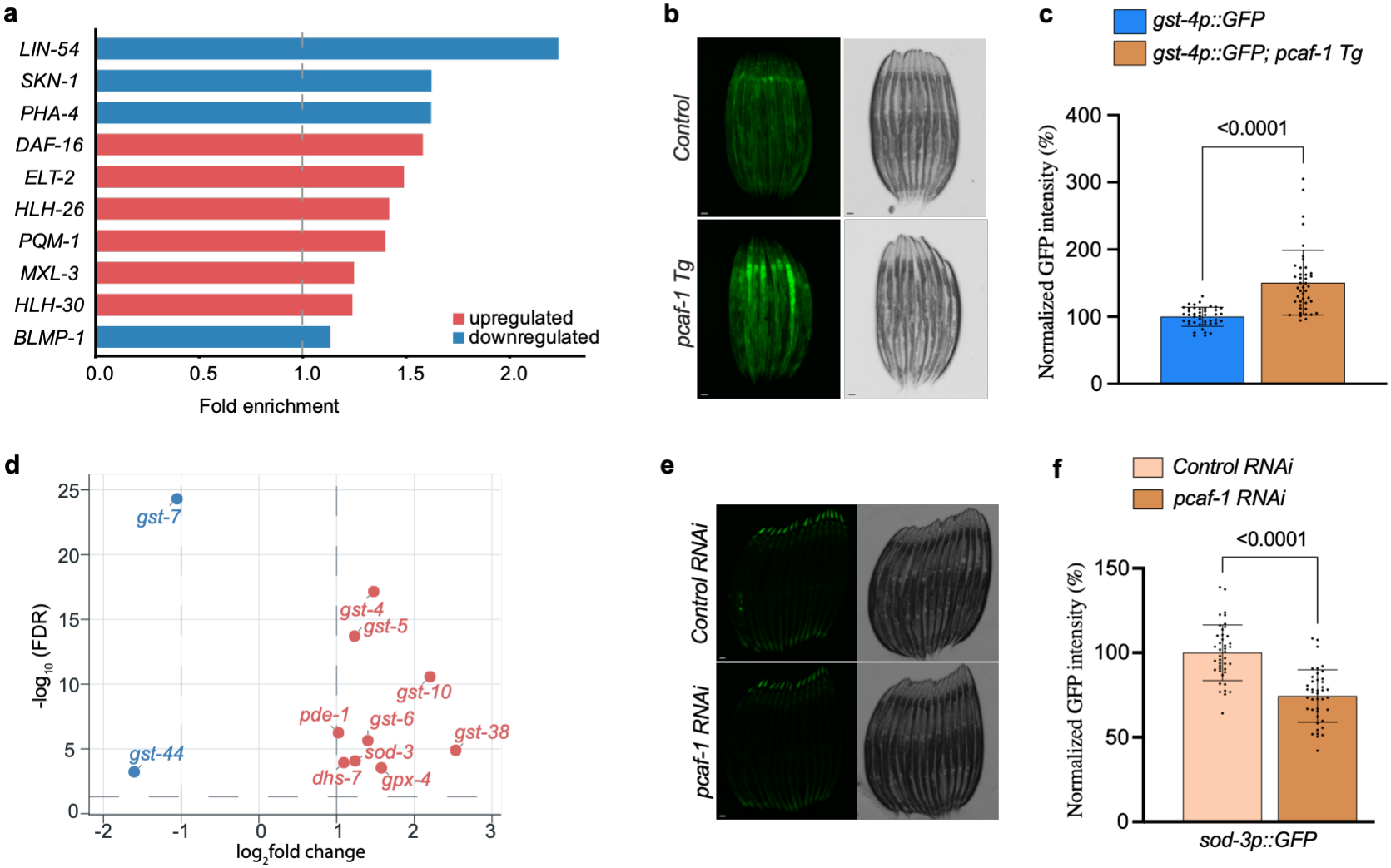
PCAF-1 regulates stress-responsive transcription and oxidative-stress-associated gene expression. **a,** WormExp enrichment of transcription-factor target signatures among genes upregulated or downregulated in *pcaf-1 Tg* animals. Red and blue bars indicate enrichment among upregulated and downregulated genes, respectively. The dashed line indicates a fold enrichment of 1. Gene lists were defined using adjusted *P* < 0.05 and |log₂ fold change| ≥ 0.263. Enrichment was assessed using the EASE-modified Fisher exact test, with Benjamini–Hochberg correction for multiple testing. **b,c,** Representative fluorescence and bright-field images (**b**) and quantification (**c**) of *gst-4p::GFP* expression in day-1 adults with or without *pcaf-1 Tg*. Approximately 14–15 animals per genotype were examined in each of three independent biological experiments. GFP fluorescence was normalized to the control mean. Bars show mean ± s.d., with individual animals shown as dots. Statistical significance was assessed using a two-sided Mann–Whitney test (*P* < 0.0001). **d,** Volcano plot of differentially expressed genes associated with reactive oxygen species metabolism, antioxidant activity and oxidant detoxification in *pcaf-1 Tg* animals. Dashed lines indicate |log₂ fold change| = 1 and FDR = 0.05. Significantly upregulated and downregulated genes are highlighted in red and blue, respectively. **e,f,** Representative fluorescence and bright-field images (**e**) and quantification (**f**) of *sod-3p::GFP* expression in day-1 adults maintained on control or *pcaf-1* RNAi from hatching. Approximately 14 animals per condition were examined in each of three independent biological experiments. GFP fluorescence was normalized to the control RNAi mean. Bars show mean ± s.d., with individual animals shown as dots. Statistical significance was assessed using a two-sided Mann–Whitney test (*P* < 0.0001). Scale bars, 50 µm (b,e).

**Extended Data Fig. 9.**
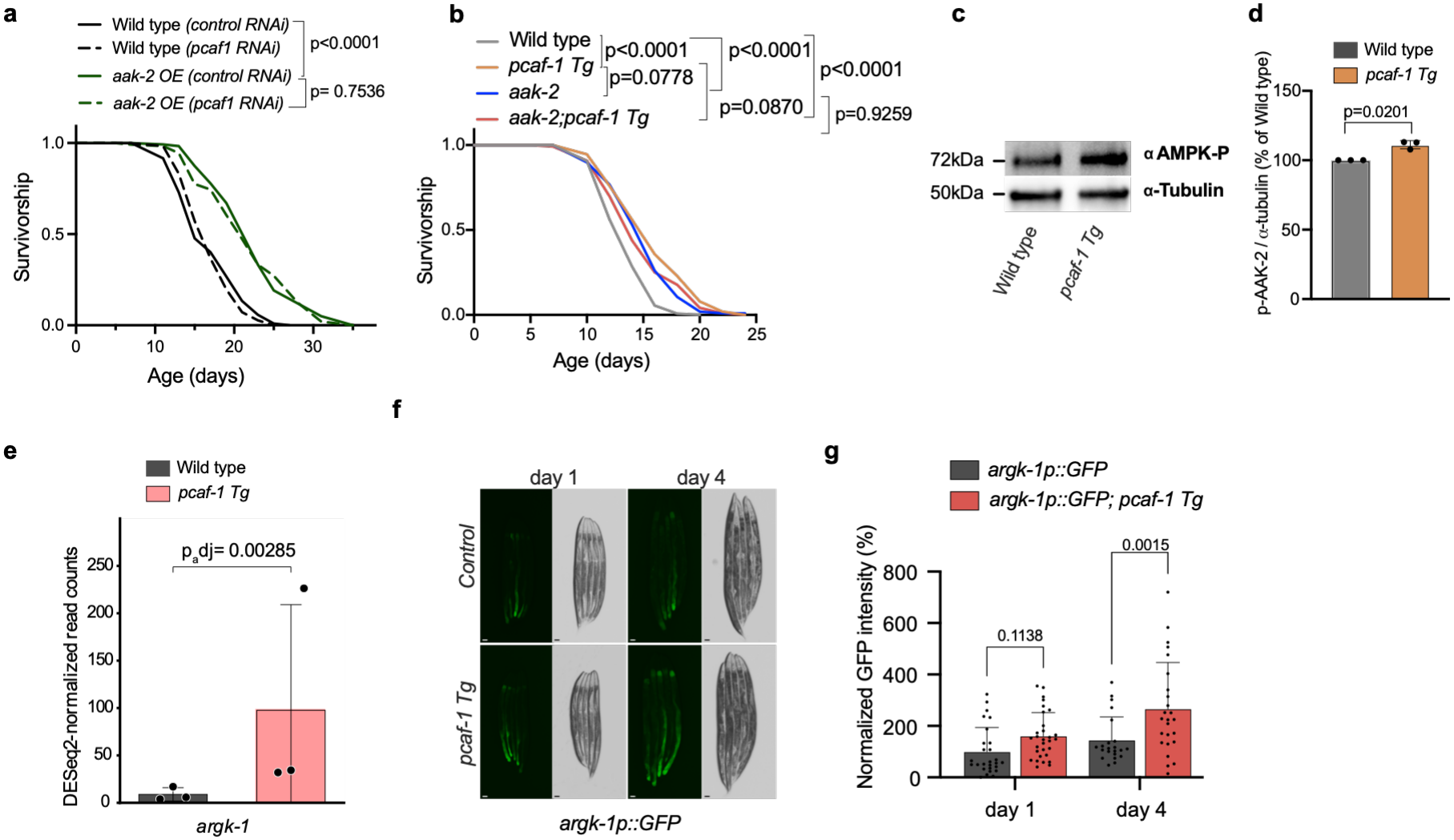
Increased PCAF-1 dosage is associated with AAK-2/AMPK signaling and *argk-1* induction. **a,** Lifespan of wild-type and AAK-2(1–321)-expressing animals maintained on control or *pcaf-1* RNAi from hatching. Survival curves were compared using the log-rank (Mantel–Cox) test. *pcaf-1* RNAi did not significantly alter the lifespan of AAK-2(1–321)-expressing animals (*P* = 0.7536). **b,** Lifespan of wild-type, *pcaf-1 Tg*, *aak-2* mutant and *aak-2; pcaf-1 Tg* animals. *pcaf-1 Tg* increased median lifespan from 14 to 16 days in the wild-type background, whereas *aak-2; pcaf-1 Tg* animals had a median lifespan of 14 days. The difference between the *pcaf-1 Tg* and *aak-2; pcaf-1 Tg* survival curves did not reach statistical significance (*P* = 0.0870). Survival curves were compared using the log-rank (Mantel–Cox) test; additional pairwise comparisons are indicated in the figure. **c,d,** Representative western blot (**c**) and quantification (**d**) of phosphorylated AAK-2 (p-AAK-2) detected using an antibody against mammalian phospho-AMPKα Thr172 and α-tubulin in day-4 wild-type and *pcaf-1 Tg* animals under fed conditions. α-Tubulin was used as a loading control. p-AAK-2 abundance was normalized to α-tubulin and expressed relative to the matched wild-type control. Data are mean ± s.d. from three independent biological experiments. Statistical significance was assessed using a two-sided one-sample *t*-test on log₂-transformed *pcaf-1 Tg*/wild-type ratios (*P* = 0.0201). **e,** DESeq2-normalized RNA-seq read counts for *argk-1* in wild-type and *pcaf-1 Tg* animals. Data are mean ± s.d. from three independent biological replicates, with individual replicates shown as dots. The adjusted *P* value was calculated using the DESeq2 Wald test with Benjamini–Hochberg correction (adjusted *P* = 0.00285). **f,g,** Representative fluorescence and bright-field images (**f**) and quantification (**g**) of *argk-1p::GFP* fluorescence in control and *pcaf-1 Tg* animals on days 1 and 4 of adulthood. GFP fluorescence was measured in individual animals and normalized to the mean fluorescence of day-1 controls. Data were pooled from two independent biological experiments and are presented as mean ± s.d., with individual animals shown as dots (day 1: *n* = 27 control and 30 *pcaf-1 Tg*; day 4: *n* = 22 control and 25 *pcaf-1 Tg*). Statistical significance was assessed using ordinary two-way ANOVA followed by Šídák’s multiple-comparisons test (day 1, adjusted *P* = 0.1138; day 4, adjusted *P* = 0.0015).

