## Extended Data Figure 2. Increased PCAF-1 dosage alters the expression of protein kinase-associated genes for "The KAT2 Acetyltransferase PCAF-1 Couples Stress-Responsive Transcription to Proteostasis and Longevity"

Extended Data Fig. 2a

PCAF1 activated protein kinases

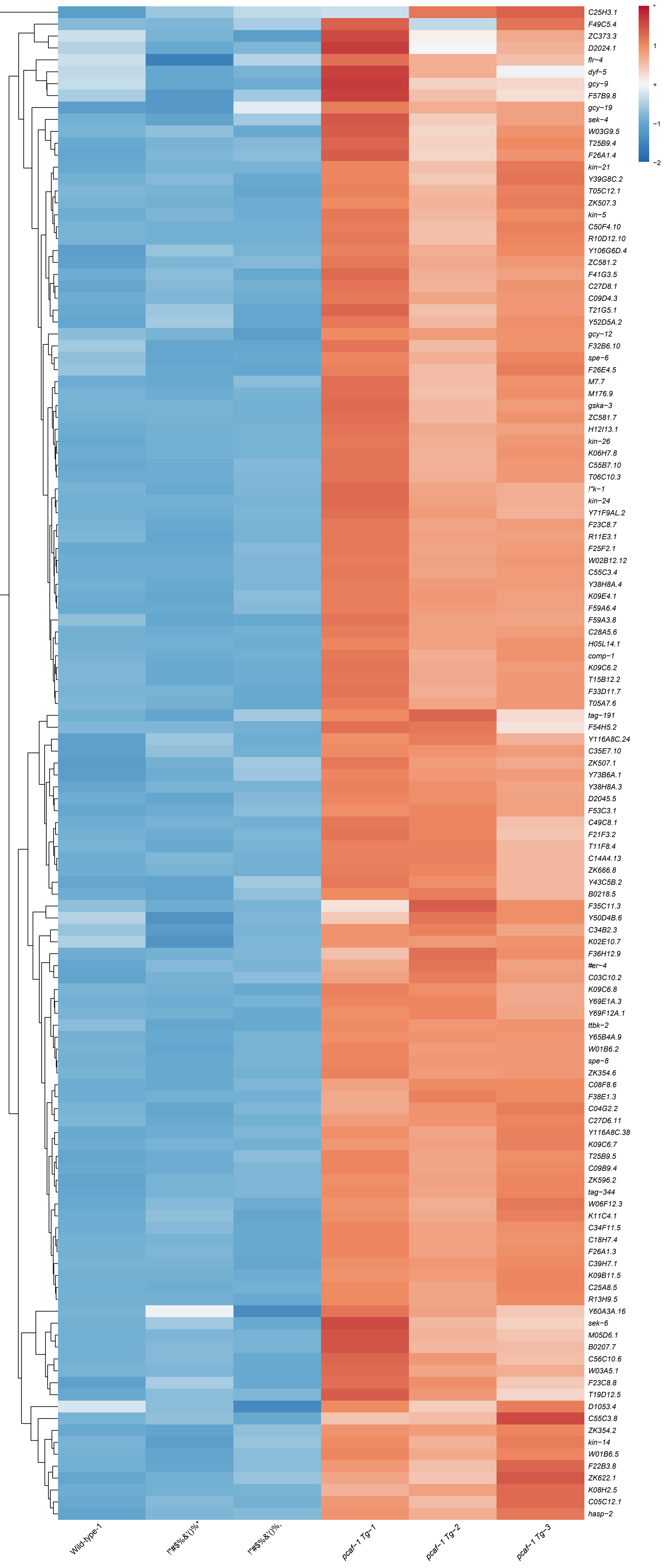

Extended Data Fig. 2b

PCAF-1 repressed protein kinases

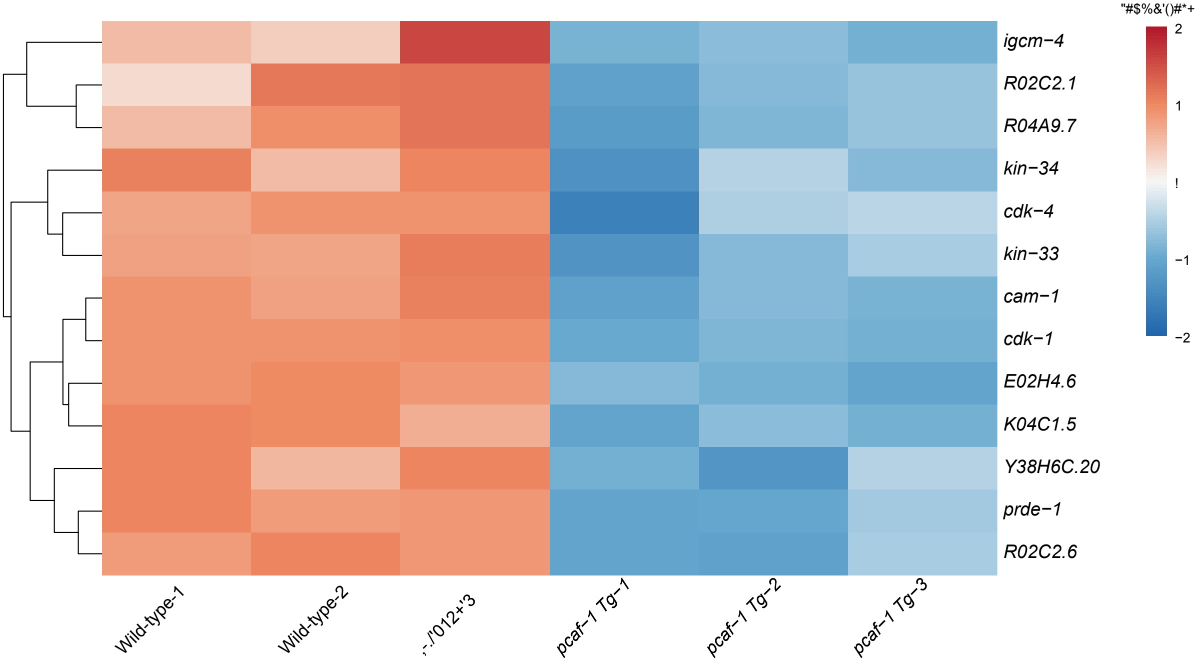
