## Extended Data Figure 4. Multiple-sequence alignment of GCN5/PCAF/KAT2-family proteins for "The KAT2 Acetyltransferase PCAF-1 Couples Stress-Responsive Transcription to Proteostasis and Longevity"

Extended Data Fig. 4

| Species | Protein | UniProt accession | Length (aa) |
| --- | --- | --- | --- |
| *Caenorhabditis elegans* | PCAF-1 | Q9N3S7 | 767 |
| *Saccharomyces cerevisiae* | GCN5 | Q03330 | 439 |
| *Drosophila melanogaster* | GCN5 | O76216 | 813 |
| *Homo sapiens* | KAT2A | Q92830 | 837 |
| *Danio rerio* | KAT2B | Q1LUC3 | 796 |
| *Mus musculus* | KAT2B | Q9JHD1 | 813 |
| *Homo sapiens* | KAT2B | Q92831 | 832 |

Alignment: MAFFT v7.526, E-INS-i. Blue shading marks columns identical in all seven proteins.

Consensus: * identical; : strongly similar; . weakly similar

Alignment columns 1-60

```
C. elegans PCAF-1        MKQRDSL-----------------------------------------------------   7
S. cerevisiae GCN5       M-----------------------------------------------------------   1
D. melanogaster GCN5     MSGGPSIT---IKSQPIDGNNTGNAAAQQQ-------------QQAAN--GAATAGASGA  42
H. sapiens KAT2A         MAEPSQAPTPAPAAQPRPLQSPAPAPTPTPAPSPASAPIPTPTPAPAPAPAAAPAGSTGT  60
D. rerio KAT2B           MSESTGI------------------------------------PQGSP--AVGAAGSAPA  22
M. musculus KAT2B        MAEAGGAG-------------------------SPA--LPPAPPHGSPRTLATAAGSSAS  33
H. sapiens KAT2B         MSEAGGAGPGGCGAGAGAGAGPGALP-----PQPAA--LPPAPPQGSP--CAAAAGGSGA  51
                         *
```

Alignment columns 61-120

```
C. elegans PCAF-1        --------------------------------------------DEKRRQLTYFTSCNA-  22
S. cerevisiae GCN5       ------------------------------------------------------------   1
D. melanogaster GCN5     AGSAQNPGHGGAASGAGSVPAEGTRQNSLQRIQQRKQKVFNLPVPQKLAKLSMYSACQS- 101
H. sapiens KAT2A         GG----PGVGSGGAGSGGDPARPG-LSQQQRASQRKAQVRGLPRAKKLEKLGVFSACKAN 115
D. rerio KAT2B           ---------APGVGGTECSGAAVG----SARIAVKKAQLRSSPRPKKLEKLGVYSSCKAE  69
M. musculus KAT2B        CG----PATAVAAAGTAEGPGGGG----SARIAVKKAQLRSAPRAKKLEKLGVYSACKAE  85
H. sapiens KAT2B         CG----PATAVAAAGTAEGPGGGG----SARIAVKKAQLRSAPRAKKLEKLGVYSACKAE 103
```

Alignment columns 121-180

```
C. elegans PCAF-1        GTCQCRGFRPVLNHEDETVERRATRAERENDFAPMTRDLETLGSEEIIECR--TCGHSIS  80
S. cerevisiae GCN5       -----------------------------------------------------VTKHQIE   8
D. melanogaster GCN5     EGCRCTGWKTPQ-------ENRHRDVE------------SSYCPEFNEECRNTSCRHSLR 142
H. sapiens KAT2A         ETCKCNGWKNPK-------PPTAPRMD-----------LQQPAANLSELCR--SCEHPLA 155
D. rerio KAT2B           GACKCNGWKSQN-PPPTPPPPTPPRAE------------QPTAVSLMEPCR--SCSHALG 114
M. musculus KAT2B        ESCKCNGWKNPN------PSPTPPRGD-----------LQQIIVSLTESCR--SCSHALA 126
H. sapiens KAT2B         ESCKCNGWKNPN------PSPTPPRAD-----------LQQIIVSLTESCR--SCSHALA 144
                                                                                * :
```

Alignment columns 181-240

```
C. elegans PCAF-1        FHAEPLKYLSQEDLDTYTDQISDLQTCCSELSEIDPSHESLQTLYITMQILLKSL--RTL 138
S. cerevisiae GCN5       E--DHLD-----------GATTDPE-----VKRVK-LENNVEEIQ---------------  34
D. melanogaster GCN5     SHIAHLDNISSSSMNELLGAIIDMENLFMSMQRVE-DEDTKKVYQYLFRLLRQCVLTRQQ 201
H. sapiens KAT2A         DHVSHLENVSEDEINRLLGMVVDVENLFMSVHKEE-DTDTKQVYFYLFKLLRKCILQMTR 214
D. rerio KAT2B           DHVTHLENVSEEEMNRLLGIVLDVEYLYTCVHKEE-DPDTKQVYFSLFKLLRKCILQMGR 173
M. musculus KAT2B        AHVSHLENVSEEEMDRLLGIVLDVEYLFTCVHKEE-DADTKQVYFYLFKLLRKSILQRGK 185
H. sapiens KAT2B         AHVSHLENVSEEEMNRLLGIVLDVEYLFTCVHKEE-DADTKQVYFYLFKLLRKSILQRGK 203
                              *.           .   * :     : . .   :. :
```

Alignment columns 241-300

```
C. elegans PCAF-1       TPVDVPFIGKPNFEPESLSPYILVKKFIASRPVDNAEKQRLAEIGTLFLTEMNHWKLEGF   198
S. cerevisiae GCN5      -----------------------------------PE-----------------------    36
D. melanogaster GCN5    AVIRGPL-GDPPFETPCITKAVLSLVFYKYNHLSTPELQTMTEVAKTFLNFLNHYNFESP   260
H. sapiens KAT2A        PVVEGSL-GSPPFEKPNIEQGVLNFVQYKFSHLAPRERQTMFELSKMFLLCLNYWKLETP   273
D. rerio KAT2B          PVVEAL--ESPPFEKPSIEQGVNNFVQYKFSHLPSKERQTIVELAKMFLNQINYWQLETP   231
M. musculus KAT2B       PVVEGSLEKKPPFEKPSIEQGVNNFVQYKFSHLPSKERQTTIELAKMFLNRINYWHLEAP   245
H. sapiens KAT2B        PVVEGSLEKKPPFEKPSIEQGVNNFVQYKFSHLPAKERQTIVELAKMFLNRINYWHLEAP   263
                                                            *
```

Alignment columns 301-360

```
C. elegans PCAF-1       QSLADKLNVPLEQGKYRMMYVRMDYYVNMPAKYVSFKQFQTVDIFGEKFTTTFCNYLINV   258
S. cerevisiae GCN5      -------------------------------------QAET-------------------    40
D. melanogaster GCN5    STRRGDLTHE-DASNYKINYTRWLVFCHVPAFCNSLRQCETSLVFGRTLLRTVFQCVSQQ   319
H. sapiens KAT2A        AQFRQRSQAE-DVATYKVNYTRWLCYCHVPQSCDSLPRYETTHVFGRSLLRSIFTVTRRQ   332
D. rerio KAT2B          SQKRQRAPDD-DVAGYKVNYTRWLCYCNVPQFCDSLPRYEATQIFGRIFLRSVFTIMRKQ   290
M. musculus KAT2B       SQRRLRSPND-DISGYKENYTRWLCYCNVPQFCDSLPRYETTKVFGRTLLRSVFTIMRRQ   304
H. sapiens KAT2B        SQRRLRSPND-DISGYKENYTRWLCYCNVPQFCDSLPRYETTQVFGRTLLRSVFTVMRRQ   322
                                                             : ::
```

Alignment columns 361-420

```
C. elegans PCAF-1       LNTGNYTPFGLSESD-LEENSED-LIT----FATDLRRYLETEDPQSTSTTNRRKDSIST   312
S. cerevisiae GCN5      ------------------------------------------------------------    40
D. melanogaster GCN5    LKKK-----CISERDRFPEDKRS-IITLMPKFLETLRAELLKDDSPIWDTSYRPSNS---   370
H. sapiens KAT2A        LLEK-----FRVEKDKLVPEKRTLILTHFPKFLSMLEEEIYGANSPIWESGFTMPPSEGT   387
D. rerio KAT2B          LLEQ-----ARQEKDKLPPEKRTLILTHFPKFLSMLEEEVYSHNSPIWSENFMIGLSGGQ   345
M. musculus KAT2B       LLEQ-----ARQEKDKLPLEKRTLILTHFPKFLSMLEEEVYSQNSPIWDQDFLSASSRTS   359
H. sapiens KAT2B        LLEQ-----ARQEKDKLPLEKRTLILTHFPKFLSMLEEEVYSQNSPIWDQDFLSASSRTS   377
```

Alignment columns 421-480

```
C. elegans PCAF-1       EDVILKYVKRKKPTDHSESSSPRELS--------------TSPVEEP---RRRKKEPSSS   355
S. cerevisiae GCN5      --------------------------------------NKQEGTDKENK-GKFEKETER-    60
D. melanogaster GCN5    ------FVIQQRKRNQEVASVPIGPSAASIGG------NKRTSVGEPLH-KRIKKEPTD-   416
H. sapiens KAT2A        QLVPRPASVSAAVVPSTPIFSPS------MGGGSNSSLSLDSAGAEPMP-GEKRTLPENL   440
D. rerio KAT2B          ----IPTVVSAPPVNRSLYYSSSPAPVELAGGGSVSPARKTASVLEPNPGGEKRKPAEPL   401
M. musculus KAT2B       PL-GIQTVIS-PPVTGTALFSSNSTSHEQINGGRTSPGCRGSSGLEANP-GEKRKMNNSH   416
H. sapiens KAT2B        QL-GIQTVINPPPVAGTISYNSTSSSLEQPNAGSSSPACKASSGLEANP-GEKRKMTDSH   435
                                                                     :     . ..
```

Alignment columns 481-540

```
C. elegans PCAF-1          A-KKSPETEESMALLRSLI-RAIKFEGDLDAMEPKTKVEFEA--LNTEVSRGLSALQEEE   411
S. cerevisiae GCN5         ----IGGSEVVTDVEKGIVKFEFD-----GVEYTFKERPSVV---------------EEN    96
D. melanogaster GCN5       ----RPSSENLDDLPADVVMRAMK-----SVSESKTTNKAEI-LFPVNVSRDENVKAEEQ   466
H. sapiens KAT2A           TLEDAKRLRVMGDIPMELVNEVML-----TITDPAAMLGPETSLLSANAARDETARLEER   495
D. rerio KAT2B             SHEDSKRPRVVGDIPMELINEVMS-----TITDPTAMLGPETSLLSAHSARDEAARLEER   456
M. musculus KAT2B          APEEAKRSRVMGDIPVELINEVMS-----TITDPAGMLGPETNFLSAHSARDEAARLEER   471
H. sapiens KAT2B           VLEEAKKPRVMGDIPMELINEVMS-----TITDPAAMLGPETNFLSAHSARDEAARLEER   490
                                   .    :   ::   :          .                       **.
```

Alignment columns 541-600

```
C. elegans PCAF-1          SGLIEFRVIGNDLDPFQHHEQLVHLVELQNLFGAQLPKMPKEYVTRLIFDSRHQNMVILK   471
S. cerevisiae GCN5         EGKIEFRVVNNDNT----KENMMVLTGLKNIFQKQLPKMPKEYIARLVYDRSHLSMAVIR   152
D. melanogaster GCN5       KRAIEFHVVGNSLTKPVDKQTVLWLLGLQLVFAYQLPEMPREYISQLVFDTKHKTLALIK   526
H. sapiens KAT2A           RGIIEFHVIGNSLTPKANRRVLLWLVGLQNVFSHQLPRMPKEYIARLVFDPKHKTLALIK   555
D. rerio KAT2B             RGVIEFHVIGNSLNQKPNKKILMWLVGLQNVFSHQLPRMPKEYITRLVFDPKHKTLSLIK   516
M. musculus KAT2B          RGVIEFHVVGNSLNQKPNKKILMWLVGLQNVFSHQLPRMPKEYITRLVFDPKHKTLALIK   531
H. sapiens KAT2B           RGVIEFHVVGNSLNQKPNKKILMWLVGLQNVFSHQLPRMPKEYITRLVFDPKHKTLALIK   550
                              ***:*:.*.      :. :: *  *: :*  ***.**:**:::*::*  * .: :::
```

Alignment columns 601-660

```
C. elegans PCAF-1          RDMGVIGGICFRTFPSRGFVEIVFCAITAMEQVKGYGTHLMNHCKDYMIK-NKIYHMLTY   530
S. cerevisiae GCN5         KPLTVVGGITYRPFDKREFAEIVFCAISSTEQVRGYGAHLMNHLKDYVRNTSNIKYFLTY   212
D. melanogaster GCN5       -ENQPIGGICFRPFPSQGFTEIVFCAVTMSEQVKGYGTHLMNHLKDYSIQ-RGIKHLLTF   584
H. sapiens KAT2A           -DGRVIGGICFRMFPTQGFTEIVFCAVTSNEQVKGYGTHLMNHLKEYHIK-HNILYFLTY   613
D. rerio KAT2B             -DGRVIGGICFRMFPTQGFTEIVFCAVTSNEQVKGYGTHLMNHLKEYHIK-HEILNFLTY   574
M. musculus KAT2B          -DGRVIGGICFRMFPSQGFTEIVFCAVTSNEQVKGYGTHLMNHLKEYHIK-HEILNFLTY   589
H. sapiens KAT2B           -DGRVIGGICFRMFPSQGFTEIVFCAVTSNEQVKGYGTHLMNHLKEYHIK-HDILNFLTY   608
                                :*** :* * .: *.******::  ***:***:***** *:*  :   *  :**:
```

Alignment columns 661-720

```
C. elegans PCAF-1          ADEFAIGYFTKQGFSEKLEINDTVYQGWIKEYEGATLMGCHLHPQISYTKFPDFSKGIQA   590
S. cerevisiae GCN5         ADNYAIGYFKKQGFTKEITLDKSIWMGYIKDYEGGTLMQCSMLPRIRYLDAGKILLLQEA   272
D. melanogaster GCN5       ADCDAIGYFKKQGFSKDVKLARPVYAGYIKEYDSATLMHCELHPSIVNTQFIAVIRSQSE   644
H. sapiens KAT2A           ADEYAIGYFKKQGFSKDIKVPKSRYLGYIKDYEGATLMECELNPRIPYTELSHIIKKQKE   673
D. rerio KAT2B             ADEYAIGYFKKQGFSKDIKVPKSKYVGYIKDYEGATLMGCELNPCIPYTEFSVIIKKQKE   634
M. musculus KAT2B          ADEYAIGYFKKQGFSKEIKIPKTKYVGYIKDYEGATLMGCELNPQIPYTEFSVIIKKQKE   649
H. sapiens KAT2B           ADEYAIGYFKKQGFSKEIKIPKTKYVGYIKDYEGATLMGCELNPRIPYTEFSVIIKKQKE   668
                           **  *****.****::.: :  . : *:**:*:..*** * : * *   .   .    .
```

Alignment columns 721-780

```
C. elegans PCAF-1            LHCGYKSENGAESRGKVFGGLEHLFRE--SSPQLLELRKVPGTDSLKMHKKSC-----YH   643
S. cerevisiae GCN5           ALRRKI-RTISKSH-IVRPGLE-QFKDLNNIKP-IDPMTIPGLKEAGWTPEMD-ALAQRP   327
D. melanogaster GCN5         ILKELIAQRHNEVQ-KVRPGLT-CFKE--GLPV-IPVESIPGLREIGWKPQNRPARSSRP   699
H. sapiens KAT2A             IIKKLIERKQAQIR-KVYPGLS-CFKE--GVRQ-IPVESVPGIRETGWKPLGK-----EK   723
D. rerio KAT2B               IIKKLIERKQAQIR-KVYPGLS-CFKE--GVRQ-IAIESIPGIRETGWKPLG-------K   682
M. musculus KAT2B            IIKKLIERKQAQIR-KVYPGLS-CFKD--GVRQ-IPIESIPGIRETGWKPSGK-----EK   699
H. sapiens KAT2B             IIKKLIERKQAQIR-KVYPGLS-CFKD--GVRQ-IPIESIPGIRETGWKPSGK-----EK   718
                                    .   : :  *  **   *::  .    :   .:**  .
```

Alignment columns 781-840

```
C. elegans PCAF-1            LDERDD--SLDSKIGAILKKLTADKNAWPFASPVDVKEVPEYYDHIKHPIDFKTMQEKLK   701
S. cerevisiae GCN5           KRGPHD-----AAIQNILTELQNHAAAWPFLQPVNKEEVPDYYDFIKEPMDLSTMEIKLE   382
D. melanogaster GCN5         LEESTDPEKLATSFASVLQSVRQHTTAWPFLRPVTAAEVPDYYDHIKYPMDLKTMGERLK   759
H. sapiens KAT2A             GKELKDPDQLYTTLKNLLAQIKSHPSAWPFMEPVKKSEAPDYYEVIRFPIDLKTMTERLR   783
D. rerio KAT2B               SKELKDPDQLYSTLKNILTQVKSHPNAWPFMEPVKKNEAPGYYQVIRFPMDLKTMSERLK   742
M. musculus KAT2B            SKEPKDPEQLYSTLKNILQQVKNHPNAWPFMEPVKRTEAPGYYEVIRFPMDLKTMSERLR   759
H. sapiens KAT2B             SKEPRDPDQLYSTLKSILQQVKSHQSAWPFMEPVKRTEAPGYYEVIRFPMDLKTMSERLK   778
                                  *     : :  :* .:  .  ****  **   *.* **: *: *:*:.**  :*.
```

Alignment columns 841-900

```
C. elegans PCAF-1            RKAYTHQHLFIADLNRLFQNCYVFNGAEAVYYKYGYKLNELALKLLKTSFPESSWYPELP   761
S. cerevisiae GCN5           SNKYQKMEDFIYDARLVFNNCRMYNGENTSYYKYANRLEKFFNNKVK----------EIP   432
D. melanogaster GCN5         KGYYQTRRLFMADMARIFSNCRFYNSPDTEYYRCANSLERYFQTKMR----------E--   807
H. sapiens KAT2A             SRYYVTRKLFVADLQRVIANCREYNPPDSEYCRCASALEKFFYFKLK----------E--   831
D. rerio KAT2B               SRYYTTRKLFMADMQRIFTNCREYNPPESEYYKCANLLEKFFYTKIK----------E--   790
M. musculus KAT2B            NRYYVSKKLFMADLQRVFTNCKEYNPPESEYYKCASILEKFFFSKIK----------E--   807
H. sapiens KAT2B             NRYYVSKKLFMADLQRVFTNCKEYNPPESEYYKCANILEKFFFSKIK----------E--   826
                                "   *   . *:  *   :: **  :*  :: * : .  *:.     ::          *"
```

Alignment columns 901-908

```
C. elegans PCAF-1            EQKPLC--                                                       767
S. cerevisiae GCN5           EYSHLID-                                                       439
D. melanogaster GCN5         --LGLWDK                                                       813
H. sapiens KAT2A             --GGLIDK                                                       837
D. rerio KAT2B               --AGLIDK                                                       796
M. musculus KAT2B            --AGLIDK                                                       813
H. sapiens KAT2B             --AGLIDK                                                       832
                                 *
```
